# Lead optimisation of dehydroemetine for repositioned use in malaria

**DOI:** 10.1101/538413

**Authors:** Priyanka Panwar, Kepa K. Burusco, Muna Abubaker, Holly Matthews, Andrey Gutnov, Elena Fernández-Álvaro, Richard A. Bryce, James Wilkinson, Niroshini Nirmalan

**Affiliations:** Environment and Life Sciences, University of Salford, Greater Manchester, M5 4WT, UK; Division of Pharmacy and Optometry, School of Health Sciences, Faculty of Biology, Medicine and Health, University of Manchester, Manchester Academic Health Science Centre, Oxford Road, Manchester M13 9PL, UK; Keele University, Staffordshire, ST5 5BG, UK; Chiroblock GMBH, Andresenstrasse 1a, 06766 Wolfen, Germany; GlaxoSmithKline, Diseases of the Developing World Medicines Development Campus, Severo Ochoa, 2. Tres Cantos, 28760, Spain

**Keywords:** malaria, antimalarial drug interactions, SYBR Green flow-cytometry, emetine, dehydroemetine, drug discovery, repositioning

## Abstract

Drug repositioning offers an effective alternative to *de novo* drug design to tackle the urgent need for novel anti-malarial treatments. The anti-amoebic compound, emetine dihydrochloride, has been identified as a potent *in-vitro* inhibitor of the multi-drug resistant strain K1 of *Plasmodium falciparum* (IC_50_: 47 nM + 2.1 nM). 2,3-dehydroemetine, a synthetic analogue of emetine dihydrochloride has been claimed to have less cardiotoxic effects than emetine. The structures of two diastereoisomers of 2,3-dehydroemetine were modelled on the reported emetine binding site on cryo-EM structure 3J7A and it was found that *(-)-R,S*-dehydroemetine mimicked the bound pose of emetine more closely than *(-)-S,S*-dehydroisoemetine. *(-)-R,S*-dehydroemetine was also found to be highly potent against the multi-drug resistant K1 strain of *P. falciparum* in comparison with *(-)-S,S*-dehydroisoemetine, which loses its potency due to the change of configuration at C-1’. In addition to its effect on the asexual erythrocytic stages of *P. falciparum*, the compounds exhibited gametocidal properties with no cross-resistance against any of the multi-drug resistant strains tested. Drug interaction studies showed *(-)-R,S*-dehydroemetine to have synergistic antimalarial activity with atovaquone and proguanil. Emetine dihydrochloride, and *(-)-R,S*-dehydroemetine failed to show any inhibition of the hERG potassium channel and displayed atovoquone-like activity on the mitochondrial membrane potential.

## Author Summary

Malaria is one of the oldest diseases on earth and has taken many lives. With nearly half of the world’s population at risk, the spread of drug resistance and insecticide resistance poses a big threat to malaria control measures. With resistance emerging to the artemisinin combination therapy (ACT), the world is in dire need of a new antimalarial. Due to long time-lines for de novo drug discovery, repositioning of drugs provides a viable alternative way. Here, we followed up on the previous repositioning work done in our lab and tested a known anti-amoebic compound for repositioned use in malaria. We found, the synthetic analogue of emetine, dehydroemetine is highly potent against the K1 multi-drug resistant strain and shows gametocidal activity with no cross-resistance. Our study provides a rationale for further optimisation of dehydroemetine for use in complicated cases of malaria.

## Introduction

Malaria presents a huge burden on the economic development of endemic countries (1). In 2016, there were 216 million malaria cases reported globally, with an estimated 445,000 deaths occurring mostly amongst african children (57). The 20^th^ century witnessed the development of a range of antimalarials including quinine alternatives such as mepacrine, chloroquine and primaquine, antifolates such as sulphadoxine and pyrimethamine, and artemisinin (2) (3) (4) (5) (6) (7). However, the emergence of resistance against all known classes of anti-malarial drugs including artemisinin chemotherapy, has warranted research into the development of new drugs with novel targets against the parasite (8) (9). Drug discovery is hindered by long development timelines, high attrition rates and soaring R&D costs (10). Antimalarial chemotherapy has predominantly relied on compounds based on natural products (11). Emetine dihydrochloride hydrate, a natural product alkaloid derived from Psychotria *ipecacuanha* has been widely used for 4 decades as a frontline anti-amoebic drug, despite its emetic and cardiotoxic side effects. It was superseded by the introduction of a safer drug metronidazole by the 1970s (12). Repositioning screens carried out at the University of Salford, UK, identified emetine dihydrochloride to have potent, nanomolar *in vitro* anti-malarial efficacy in the multi-drug resistant K1 *Plasmodium falciparum* parasite strain. The comparatively significant difference in *in vitro* anti-protozoan efficacies (IC_50_ 47 + 2.1 nM in *P. falciparum* as compared to IC_50_ 26.8 + 1.27 μM in *Entamoeba histolytica*) dictate that the safety profile for its repositioned use as an antimalarial could be different (13) (14). The pleiotropic natural product drug has been reported to have anti-viral and anti-cancer properties, including recent reports of interrupting viral replication and cell entry for Zika and Ebola viruses (15). The 40S ribosomal subunit of the eukaryotic 80S ribosome was recently reported to be the site of action of emetine (16). The definition of the target binding site for emetine enables a chimeric approach to refine using rational design, a drug discovered through repositioning.

The (*R*) configuration at C-1’ and the presence of a secondary nitrogen at position 2’ are important for emetine’s biological activity (Fig 1).

**Fig 1:**
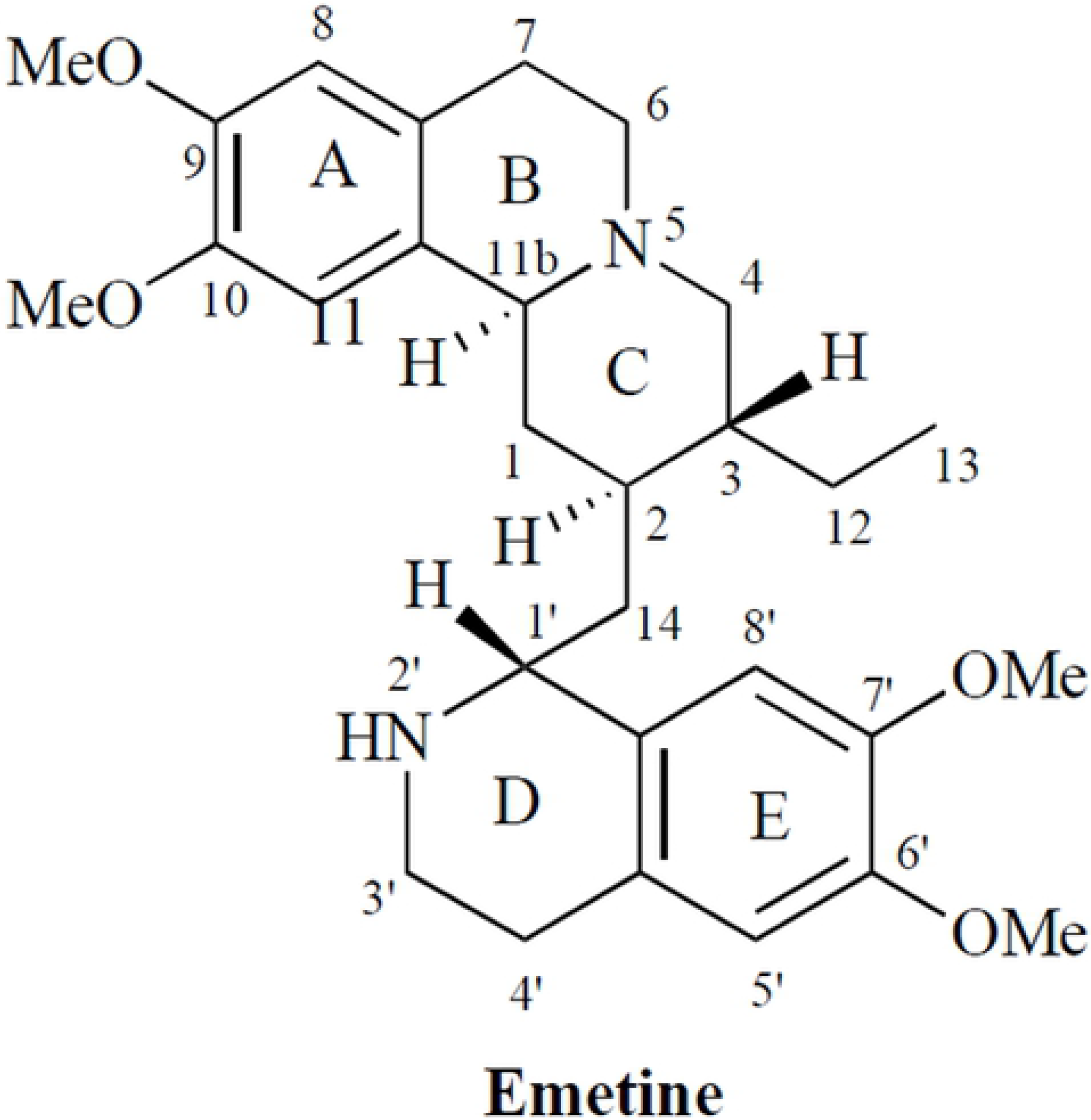
Structure of emetine hydrochloride.

Indeed, the (*S*) configuration at C-1’ (isoemetine) or the substitution of the secondary amine result in loss of activity. Even if the asymmetry at carbons 2 and 3 are lost with unsaturation at position 2-3 (2,3-dehydroemetine) the biological activity is retained (17). In 1980, a study was conducted on the cross-resistance of emetine-resistant mutants of Chinese hamster ovary cells to related compounds such as (-)-cryptopleurine (phenanthroquinolizidine-type alkaloids), (-)-tylocrebrine (phenanthroindolizidine type), and (-)-emetine, (-)-tubulosine, (-)-cephaeline, and (-)-dehydroemetine (benzoiosoquinoline alkaloids). They concluded that the planar structure of the molecule, with two aromatic rings was made slightly electronegative by hydroxyl or methoxyl groups (18). The distance between the two aromatic rings and the angle between the nucleophilic element, such as nitrogen, and the rings were essential for biological activity.

Emetine in its crude form has been in use since 1658 for the treatment of dysentery and was first brought to Europe from Brazil by Piso (19). Large doses of emetine in powdered ipecacuanha form were introduced in Mauritius in 1858 and the annual death rate from severe dysentery was reduced from 10-18% to 2% (20). Generalised muscle weakness, vomiting and cardiotoxicity were the side-effects with prolonged use of emetine, as in the treatment of hepatic amoebiasis (21) (22).The synthetic analogue of emetine, 2,3-dehydroemetine, was introduced in 1959 (23). It was found to be less emetic than emetine and could be given orally as a resinate. 2,3-Dehydroemetine was also comparatively safer than emetine when given parenterally at similar doses. At high doses, the electrocardiographic changes were similar but were less marked and of shorter duration (24). Experimental results have also shown that changes observed in heart conduction, contractility, automaticity and ECG abnormalities by emetine and 2,3-dehydroemetine could be due to their effect on membrane permeability to Na+, K+ and, Ca++ ions (25). 2,3-Dehydroemetine has been claimed to have a less cardio-toxic effects than emetine in a study that assessed the effect of chloroquine in combination with emetine hydrochloride and 2,3-dehydroemetine for the treatment of amoebic liver abscesses (26).

A study was conducted to observe the distribution and excretion of 2,3-dehydroemetine and emetine in guinea pigs (27). It was found that 91.5% of 2,3-dehydroemetine compared to 67% of emetine was excreted 72 hours after stoppage of doses. 2,3-Dehydroemetine was eliminated more rapidly from the heart than the liver and reverse was found to be true for emetine. This could be one of the reasons for reduced cardiotoxicity observed in treatment with 2,3-dehydroemetine, as emetine binds more tightly to cardiomyocytes even though it is not specifically concentrated in them. Less toxic analogues have not been analysed for antimalarial potency, therefore, based on the anecdotal evidence, two diastereoisomers of dehydroemetine, *(-)-R,S*-dehydroemetine and *(-)-S,S*-dehydroisoemetine were synthesised. Molecular modelling tools (28) (29) were used to predict the activity of the diastereoisomers and their potency against the multi-drug resistant K1 strain of *P. falciparum* was tested.

It was observed that 2,3-dehydroemetine and emetine were potent inhibitors of protein synthesis in HeLa cells (17). At a concentration of 5 × 10^−8^ M, emetine was able to effectively inhibit protein synthesis by 50% in HeLa cells but had no effect on the incorporation of amino acids into the mitochondrial fraction even at 1.4 × 10^−4^ M (30). On cultured chicken embryo cardiac cells, emetine led to cessation or progressively decreased rate of beating in atrial and ventricular cells depending on the concentration. In the study washing did not reverse the effects but the addition of NADH (10^−5^ M) resulted in prolonged reversal lasting 30 to 45 minutes whereas intense brief reversal of effects was observed by addition of ATP (10^−6^ M) indicating that at higher concentrations, emetine causes physical disruption of the plasma membrane of the cells and at a lower concentration, it suppresses the process of formation of high energy phosphate compounds (31). It was thought that the site of inhibition was at the enzymatic oxidation of substrates mediated by nicotinamide adenine dinucleotide (NAD). In this study we tested the activity of emetine dihydrochloride and its two synthetic analogues against hERG potassium channel. Drugs inhibiting the hERG channel can prolong the QT interval and cause a dangerous cardiac arrhythmia, *Torsades de pointes* (32). The three compounds were also tested for any effect on mitochondrial membrane potential.

Parasites are transmitted from the mammalian hosts to mosquito vectors through mature *Plasmodium* gametocytes and hence the reinfection cycle could be broken with the use of transmission blocking anti-malarials (33). Cross-resistance and gametocidal activity of emetine dihyrochloride and its synthesised analogues were tested by GSK in a bid to determine the potential of these compounds as transmission blocking drugs.

The preference of combinatorial regimes over monotherapy for the treatment of malaria has affected the drug discovery pipeline in a crucial way (34). More often than not, instead of developing two new drugs for combination therapy, newer drug candidates are tested for synergistic activities with existing anti-malarial treatments (35). Combination therapy not only helps in dose reduction and better therapeutic efficacy, it is also useful in delaying the emergence of resistance and widens the shelf-life of the existing cohort of drugs (36).

Chou and Talalay developed a method based on the argument that the issue of synergy is more physiochemical rather than statistical in nature and employed the mass-action law principle to derive a median-effect equation where additivity could be defined using the resulting combination index (CI = 1) with antagonism and synergism defined as >1 and <1 respectively. Calcusyn and Compusyn are two software programmes based on the complex algorithms for median-effect analysis to allow automation and eliminate subjectivity during data analysis (37). We have previously demonstrated the use of Calcusyn as a reliable method to define anti-malarial drug interactivity for combinatorial regimes (38).

Hence, to reduce dose-dependent side effects, combinatorial partner drugs showing synergistic activity were sought. *(-)-R,S*-Dehydroemetine was taken forward for drug interaction studies as it was found to be highly potent against the K1 strain of *P. falciparum*. Atovaquone and proguanil were first evaluated to test the efficacy of the method as the two drugs are known to exhibit synergism (39). The methodology was then applied to evaluate combinatorial partner drugs for the potent anti-malarial candidate (-)-*R,S*-dehydroemetine. Pre-determined IC_50_ values for each compound were used to select the constant-ratio for each combination.

## Results

### Molecular modelling of ligand interactions with *P. falciparum* 80S ribosome

In order to explore the structural basis of the relative inhibitory activity of *(-)-R,S*-dehydroemetine and *(-)-S,S*-dehydroisoemetine, we predicted and compared their molecular interaction with *P. falciparum* 80S ribosome using computational docking. As the receptor structure, we use the recently solved cryo-EM structure of the 80S ribosome of *P. falciparum*, determined to a resolution of 3.2 Å (16). The electron density corresponding to the bound emetine is located in the E-site of the ribosomal small subunit, ie. *Pf*40S; this E-site, a binding site that pactamycin also recognises in the bacterial 30S subunit, is at the interface between 18S rRNA helices 23, 24, 45 and the C-terminus of protein uS11.

On closer examination of the published cryo-EM structure of the complex, the ligand structure originally modelled into the density, unfortunately, corresponds to the non-natural enantiomer of emetine, (1*S*,2*R*,3*S*,11b*R*)-emetine. This modelled structure was not the ligand employed in the associated cryo-EM experiments (16). Therefore, we docked emetine, i.e. the (1*R*,2*S*,3*R*,11b*S*) structure, into the identified *Pf*40S binding site using MOE-Dock [Chemical Computing Group Inc., 1010 Sherbrooke St. West, Suite #910, Montreal, QC, Canada, H3A 2R7, 2016.]. The docked emetine geometry maps well into the electron density envelope (shown at a contour level 0.1542e/A^3 (3.50rmsd) in Fig 2a); the bound pose broadly follows the twisted U-shape conformation of the observed electron density, resembling some aspects of the previously modelled non-natural emetine enantiomer geometry (Fig 2b).

**Fig 2:**
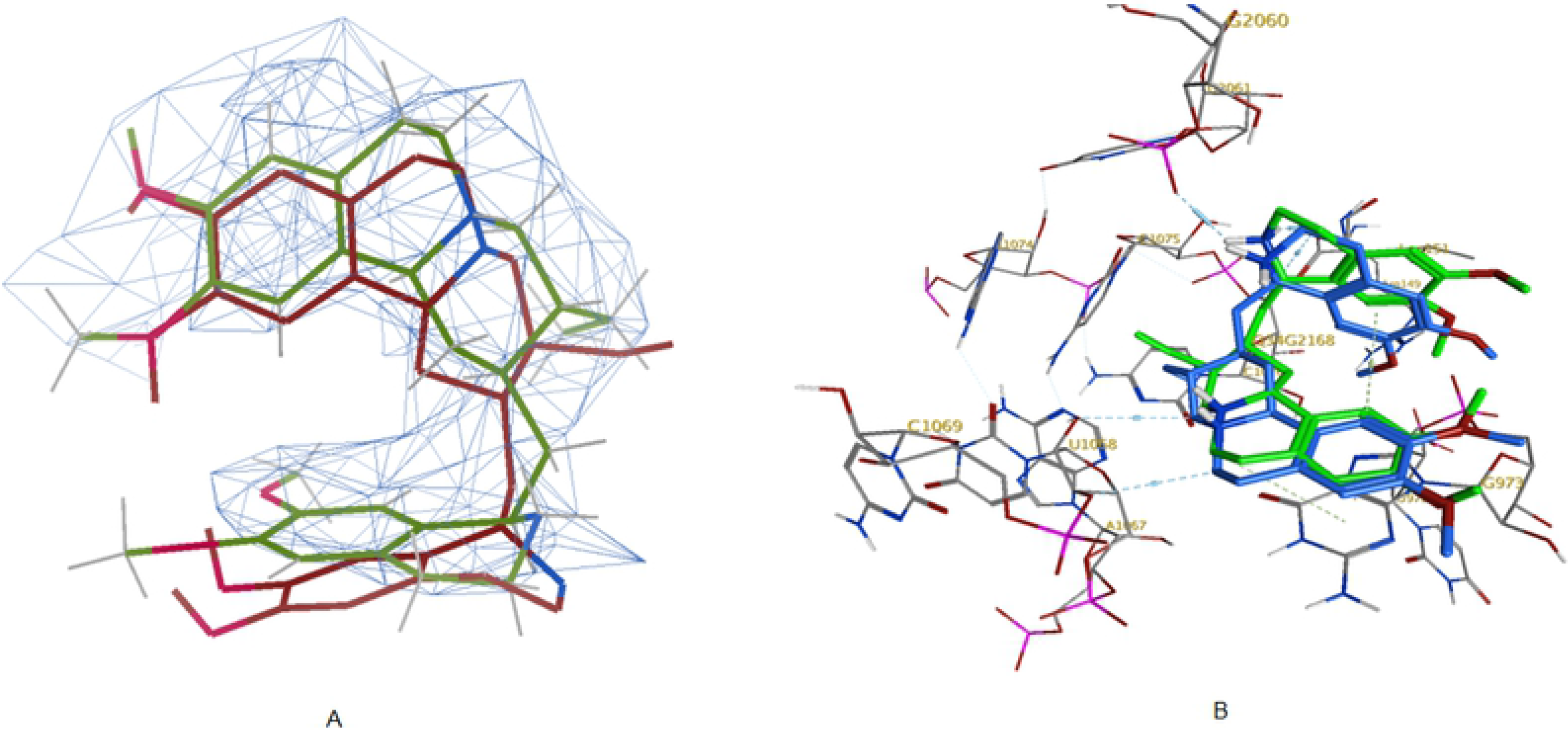
(a) Overlay of docked pose of emetine (green) with its enantiomer present in the cryo-EM structure (maroon); observed electron density envelope is also shown (wireframe surface with contour level 0.1542e/A^3 (3.50rmsd)); (b) Interactions of docked emetine (green) with Pf40S residues and comparison with previously modelled interactions of its enantiomer (maroon) (16)

In this U-shape, an intermolecular T-shaped π-stacking interaction is observed between the two cyclic systems of emetine, ie. benzo[a]quinolizine rings A/B/C and isoquinoline rings D/E (Fig 1, Fig 2a). The docked emetine also forges a number of comparable interactions with the *Pf*40S subunit as its modelled enantiomer (Fig 2b), with a key π-stacking interaction between the A/B/C rings of emetine with the purine ring of Gua973 of h23 (16). The location of the emetine secondary and tertiary amines are broadly similar (within 1 - 2 Å), allowing hydrogen bonding interactions with a backbone oxygen atom of Ura2061 (h45) and the 2’-hydroxyl group of Ura1068 (h24) respectively (Fig 2b). The tertiary amine also forms a salt bridge interaction with the carboxylate side chain of C-terminal residue Leu151 of uS11 (Fig 2b). This interaction was not highlighted in the cryo-EM study and is a consequence of modelling into the E-site the natural emetine geometry.

Subsequently, the two diastereomers of emetine, *(-)-R,S*-dehydroemetine and *(-)-S,S*-dehydroisoemetine, were docked in turn into the emetine-binding region of the *Pf*40S subunit. We found that the preferred docked pose of the *(-)-R,S*-dehydroemetine adopts the familiar U-shaped conformation, superimposing rather closely onto the bound pose of emetine (Fig 3a). The docking scores are correspondingly similar for emetine, with a London dG value of −7.2 kcal/mol, and *(-)-R,S*-dehydroemetine, which has a dG score of −7.3 kcal/mol. As observed for emetine, the *(-)-R,S*-dehydroemetine forms the π–π stacking interaction with the Gua973 pyrimidine ring and polar interactions with Ura2061, Ura1068 and Leu151 (Fig 3b).

**Fig 3:**
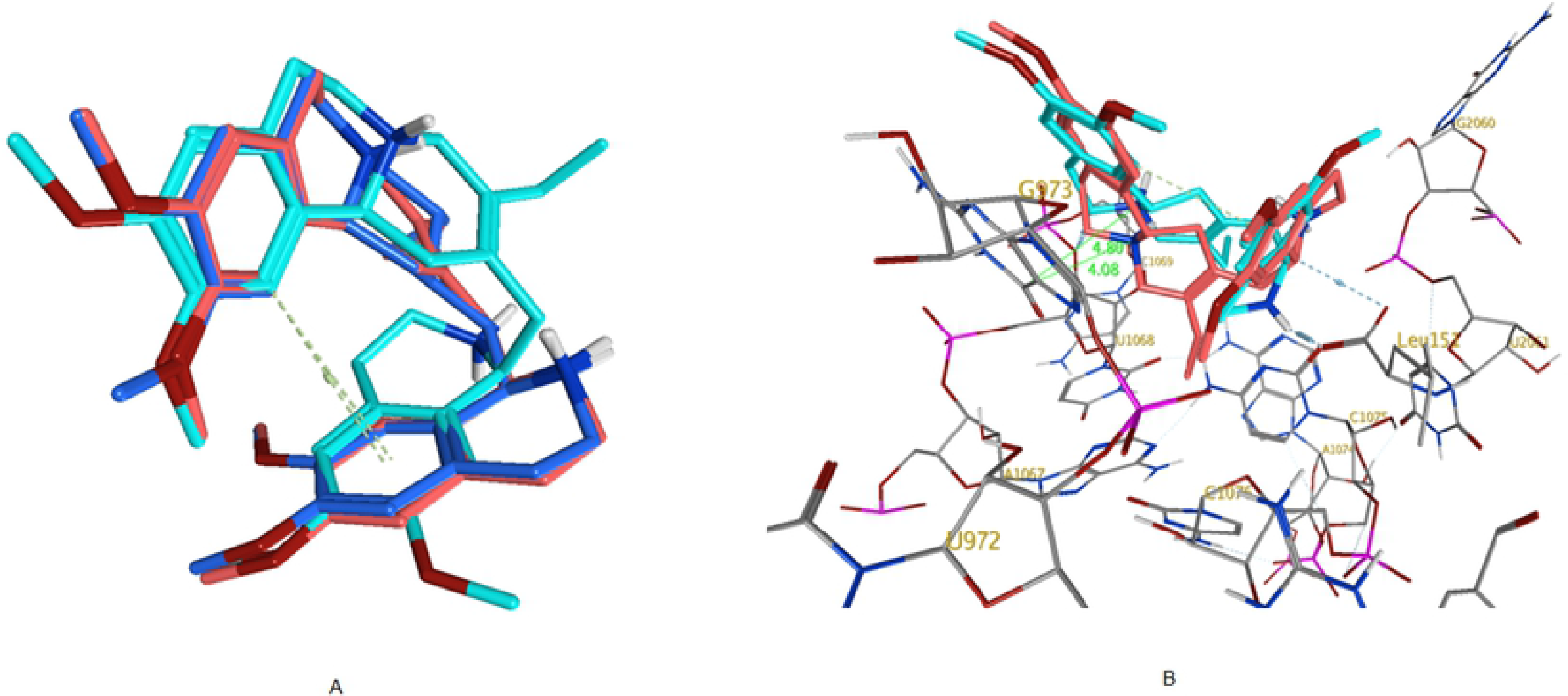
(a) Overlay of docked poses of (-)-R,S-dehydroemetine (red) and (-)-S,S-dehydroisoemetine (cyan) with emetine (blue). (b) Interactions of (-)-R,S-dehydroemetine (red) and (-)-S,S-dehydroisoemetine (cyan) with *the Pf*40S binding site. Distances (dotted lines) in Å.

However, *(-)-S,S*-dehydroisoemetine docks into the binding site with a lower dG score of −6.5 kcal/mol and does not overlay in conformation so readily with emetine or the *(-)-R,S*-dehydroemetine (Fig 3a). The particular stereochemical configuration of *(-)-S,S*-dehydroisoemetine appears to result in its secondary amine being more distant from the E-site residues of *Pf*40S. Consequently, this amine N…O Ura2061 distance extends by 0.8 Å in proceeding from R to S isomer (Fig 3b). The tertiary amine interaction with Ura1068 is maintained, however, as is the interaction with the terminal carboxylate of Leu151 (Fig 3b). The π stacking interaction with Gua973 is also present, but at a slightly larger distance between planes, increased by ~0.7 Å. Fig 4 (a) and (b) shows the predicted binding site residues for *(-)-R,S*-dehydroemetine and *(-)-S,S*-dehydroisoemetine.

**Fig 4:**
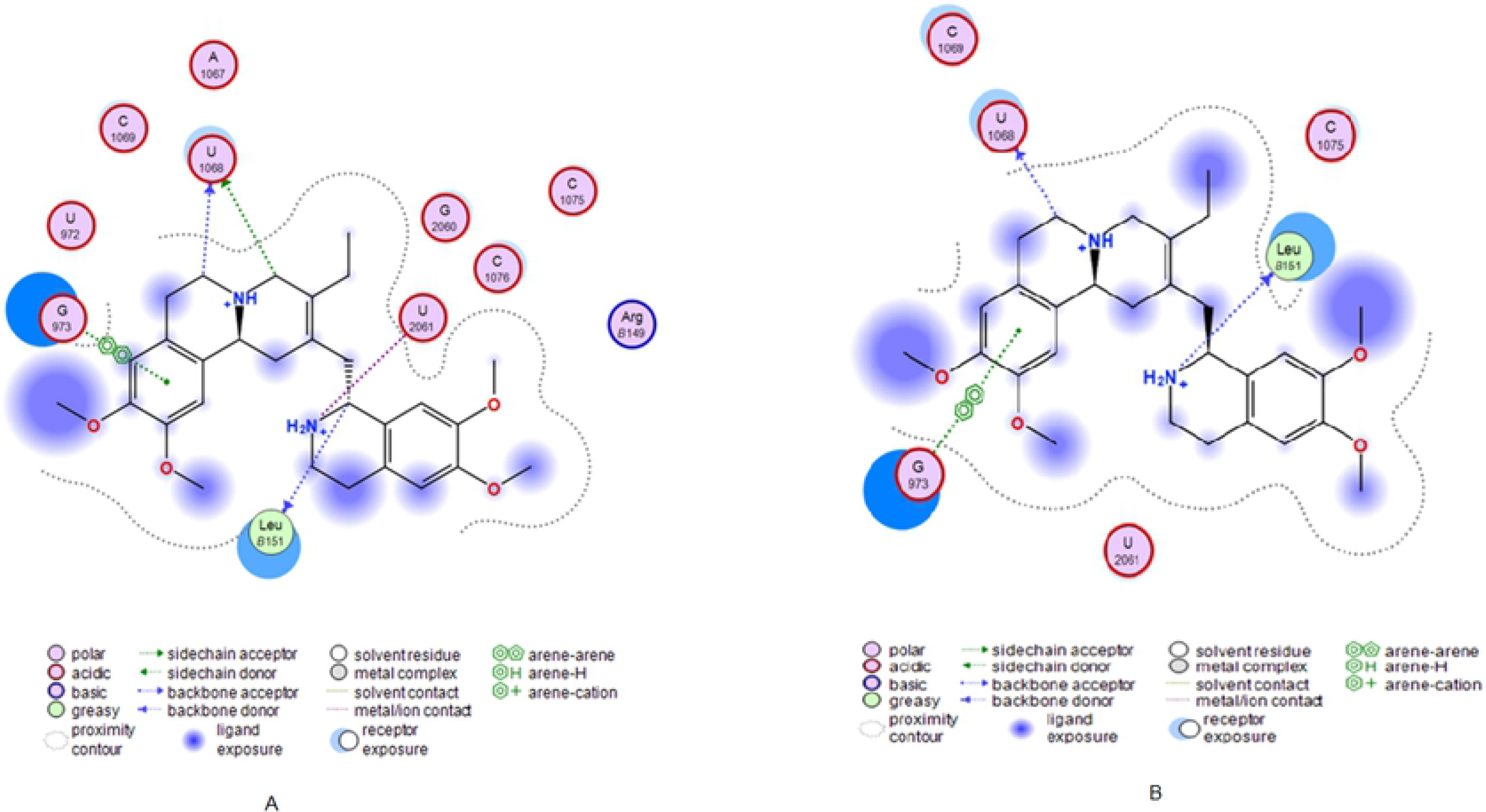
(a) Docking through MOE showing the binding site residues for (-)-R,S-dehydroemetine molecule. (b) Docking through MOE showing the binding site residues for (-)-S,S-dehydroisoemetine molecule.

### Dose-response experiments to test activity against the K1 strain of *P. falciparum*

Based on the anecdotal evidence, two synthetic analogues of emetine dihydrochloride, *(-)-R,S*-dehydroemetine and *(-)-S,S*-dehydroisoemetine were synthesised. Experiments to test drug efficacy were set up and *(-)-R,S*-dehydroemetine was tested in two fold serial dilutions from 12.5 nM to 200 nM. A dose response experiment was set up on ring stage of the K1 strain of *P. falciparum* to be read at 72 hours. IC_50_ of *(-)-R,S*-dehydroemetine was observed to be 69.58 + 2.62 nM. Results are shown in Fig 5. *(-)-S,S*-dehydroisoemetine was tested in two fold serial dilutions from 0.625 μM to 10 μM. IC_50_ was observed to be 1.85 + 0.2 μM concentration. Results are shown in Fig 5.

**Fig 5:**
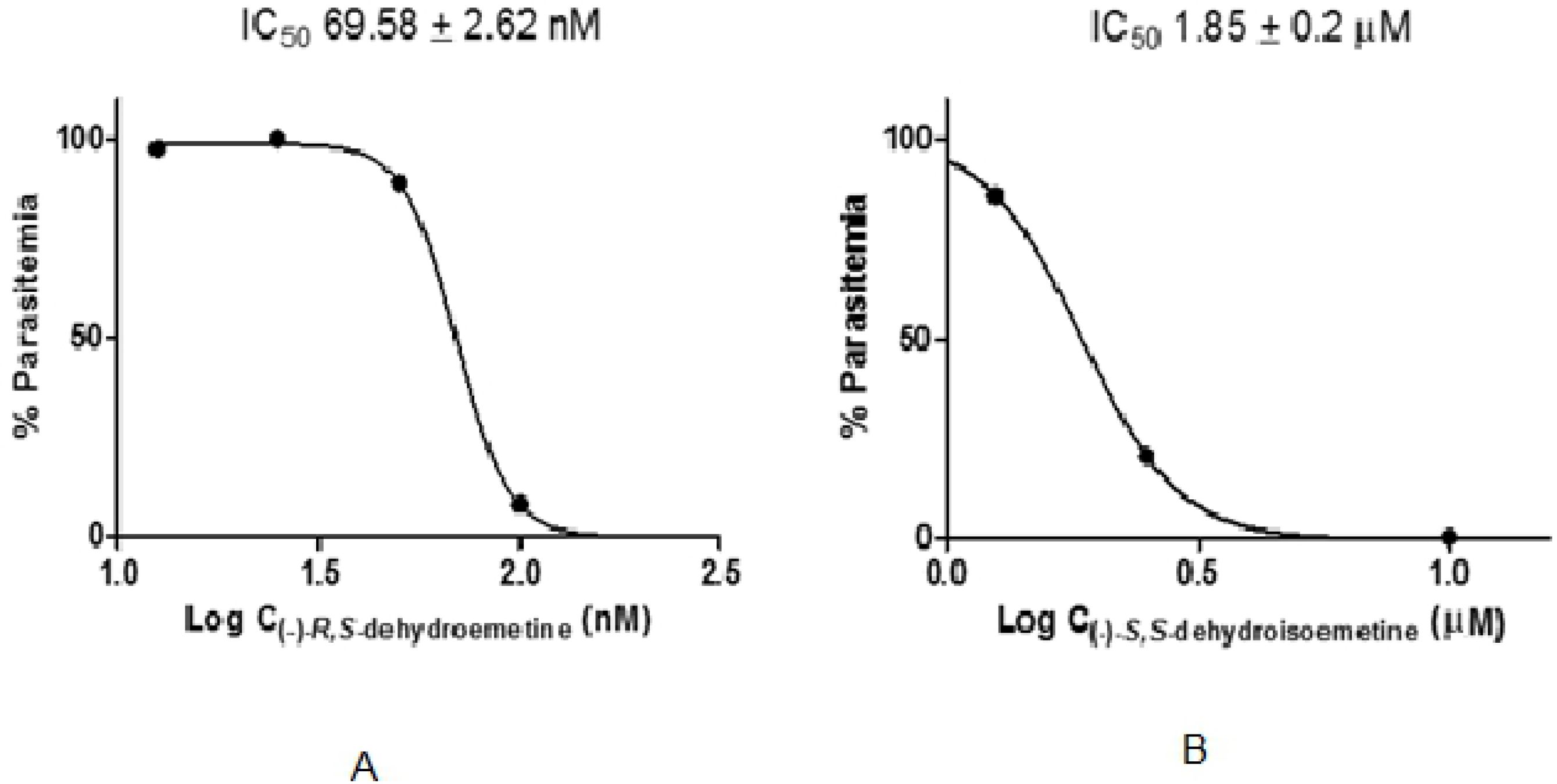
(a) Dose-response experiment for *(-)-R,S*-dehydroemetine read at 72 hours (b) Dose-response experiment for *(-)-S,S*-dehydroisoemetine read at 72 hours

### IC_50_ speed assay for *(-)-R,S*-dehydroemetine and *(-)-S,S*-dehydroisoemetine

Accurate determination of the parasite killing rate in response to treatment is crucial in the development of drugs against *P. falciparum* as chemotherapy remains the primary element in the control of malaria. The speed of action of compounds on the viability of parasites is difficult to measure through traditional techniques (40). It is important to identify new drugs with rapid parasite killing kinetics early in the drug development process. Besides rapid relief of symptoms, a fast-acting drug also helps to curtail the mutations causing the development of new mechanisms of drug resistance. In 2012, Tres Cantos developed a labour intensive low-throughput assay taking up to 28 days to determine *in vitro* the parasite reduction ratio (PRR) and the presence or absence of a lag phase in response to a drug (41). The method used in this study to differentiate between fast and slow acting compounds gives initial results in 4 to 7 days and is able to detect static or cidal effects of compounds (42).

The IC_50_ speed assay was performed for *(-)-R,S*-dehydroemetine and *(-)-S,S*-dehydroisoemetine (with concentrations of 25 nM, 125 nM, 625 nM, 3125 nM and 15625 nM and 0.5 μM, 2.5 μM, 12.5 μM, 62.5 μM and 312.5 μM, respectively) using unsynchronised cultures of *P. falciparum*. The ratio of 24 hours IC_50_ values to 72 hours IC_50_ values for *(-)-R,S*-dehydroemetine and *(-)-S,S*-dehydroisoemetine could not be determined as IC_50_ could not be reached within 24 hours. Both compounds only achieved 50% inhibition at the previously defined IC_50_ values after 48 hours of exposure, indicating that the isomers have delayed action against the multi-drug resistant K1 strain of *P. falciparum*. The results are shown in Fig 6.

**Fig 6:**
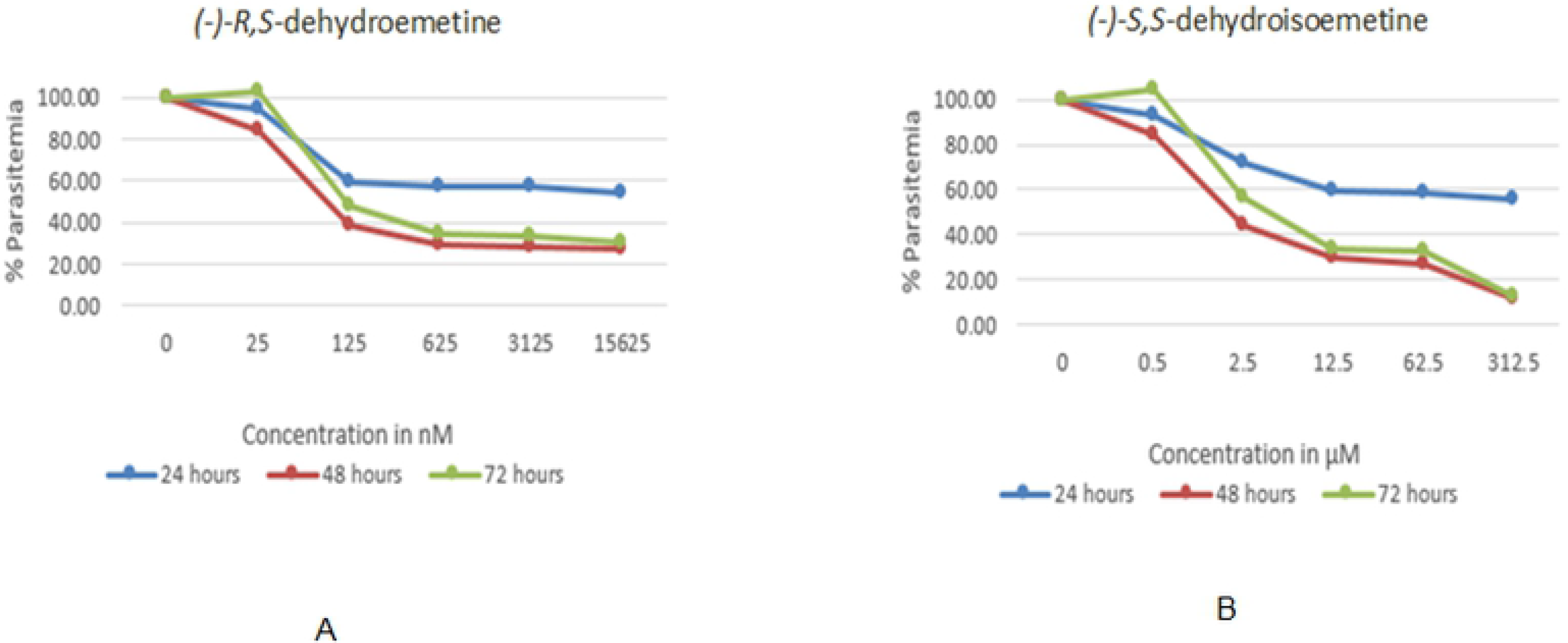
IC_50_ speed assay using unsynchronised cultures of *Plasmodium falciparum*. IC_50_ could not be reached within 24 hours. *R, S*-2,3-dehydroemetine and *(-)-S,S-dehydroisoemetine* reached the IC_50_ by 48 hours. (a) IC_50_ speed assay for *(-)-R,S*-dehydroemetine with concentration (25, 125, 625, 3125 and 15625 nM), (b) *(-)-S,S*-dehydroisoemetine with concentration (0.5, 2.5, 12.5, 62.5 and 312.5 μM)

### Stage-Specificity assay

The two isomers of 2,3-dehydroemetine were tested on synchronous cultures to determine the stage specificity of the compounds by measuring the concentration-dependent growth of schizonts and rings following incubation with the two compounds.

It was observed that *(-)-R,S*-dehydroemetine and *(-)-S,S*-dehydroisoemetine affect both the ring and trophozoite/schizont stages of the parasite. The isomers were found to be more active in the late trophozoite/schizont stages which is consistent with the proposed protein synthesis target of the parent compound. The lower potency against ring stage parasites may also in part explain the lag-phase observed during the speed assay, where the IC_50_ was not reached after 24 hours of exposure against unsynchronised cultures.

The IC_50_ speed assay suggests that both the compounds have a slow action against unsynchronised cultures. However, the stage specificity assay shows a slow activity (~40% action) against the rings even at high concentration and fast activity on late trophozoites/schizonts (~80% action) (Fig 7). By correlating the data obtained from the two assays, both *(-)-R,S*-dehydroemetine and *(-)-S,S*-dehydroisoemetine should have a slow action against the rings and fast action against the late trophozoites/schizonts stages.

**Fig 7:**
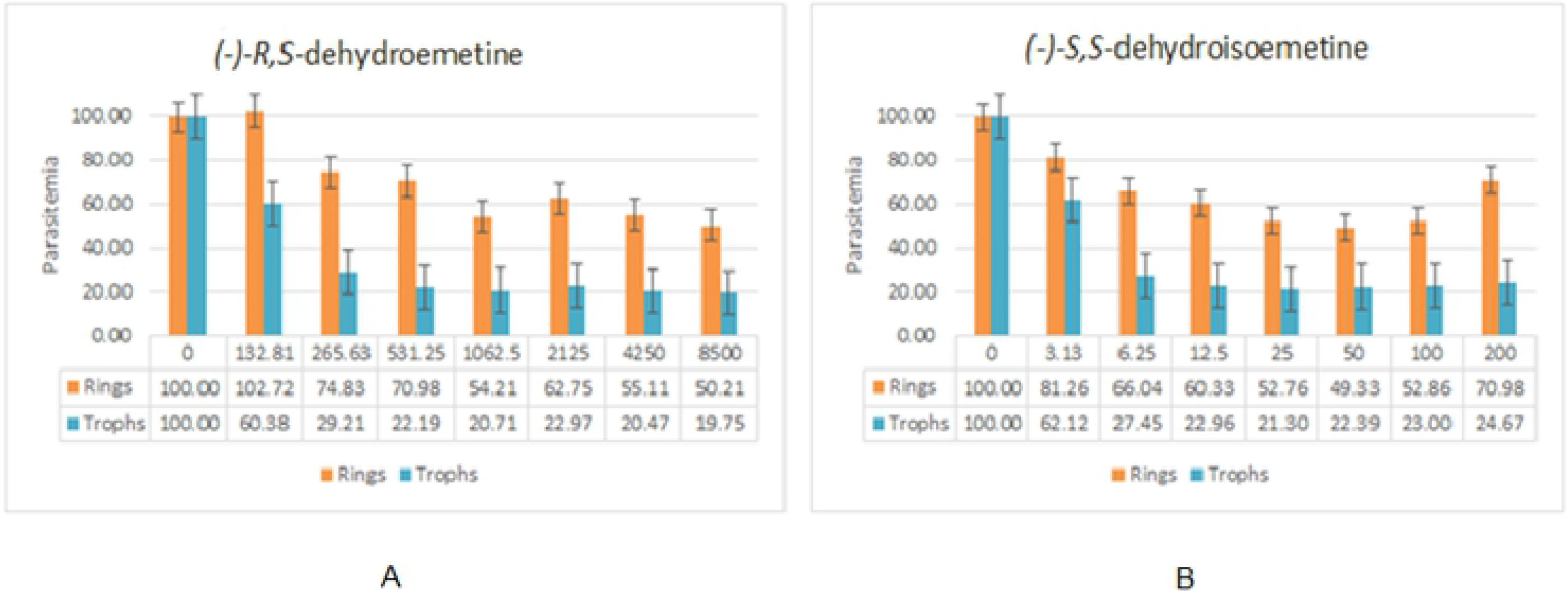
Stage specificity assay using synchronised cultures of *Plasmodium falciparum*. The decrease in growth of trophozoites/schizonts was observed to be more marked in comparison to rings thus explaining the slow action observed in the IC_50_ speed assay. (a) Stage specificity assay for *(-)-R,S*-dehydroemetine, (b) Stage specificity assay for *(-)-S,S*-dehydroisoemetine

### MTT assay for cell cytotoxicity against HepG2 cells

MTT assay for cell cytotoxicity was performed for *(-)-R,S*-dehydroemetine and *(-)-S,S*-dehydroisoemetine. Emetine and cisplatin were used as control drugs (Fig 8). The plates were read at 24 hours (43).

**Fig 8:**
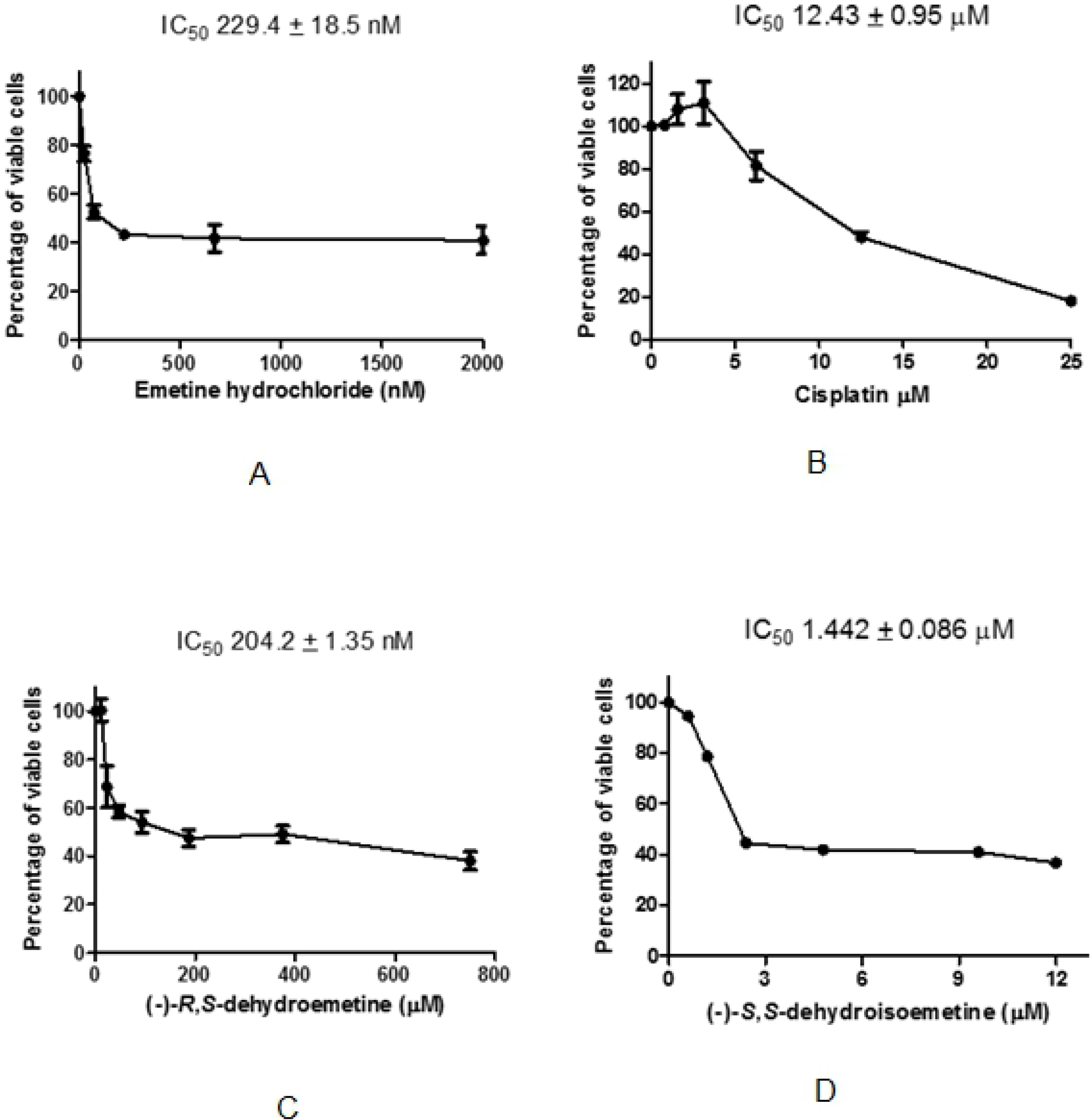
48 hour MTT assay. (a) 48 hour MTT assay for emetine, (b) 48 hour MTT assay for cisplatin, (c) 48 hour MTT assay for *(-)-R,S*-dehydroemetine and (d) 48 hour MTT assay for *(-)-S,S*-dehydroisoemetine

Emetine was tested in three-fold serial dilutions from 24.7 nM to 2000 nM with IC_50_ observed to be 229.4 + 18.5 nM. *(-)-R,S*-dehydroemetine was tested in two-fold serial dilution from 11.72 nM to 750 nM with IC_50_ observed to be 204.2 + 1.35 nM. *(-)-S,S*-dehydroisoemetine was tested in two fold serial dilutions from 0.6 μM to 12 μM with IC_50_ observed to be 1.442 + 0.086 μM. Cisplatin was tested in two fold serial dilutions from 0.78 μM to 25 μM and with IC_50_ observed to be 12.43 + 0.95 μM. The selectivity index, calculated as IC_50_ HepG2 / IC_50_ parasites (44), of *(-)-R,S*-dehydroemetine was approximately 3 and *(-)-S,S*-dehydroisoemetine were observed to be approximately equal to 1.

### Determination of cross-resistance through hypoxanthine incorporation assay

This assay relies on the parasite incorporation of labelled hypoxanthine that is proportional to *P. falciparum* growth. *In vitro* cross-resistance of the compounds is measured as an IC_50_ ratio between the IC_50_ value for the corresponding *P. falciparum* strain versus the IC_50_ value for 3D7A. Every replicate from an MDR strain involved a simultaneous determination using a 3D7A replicate to avoid any artefact linked to experimental conditions. Table 1 shows the IC_50_ values of emetine dihydrochloride, *(-)-R,S*-dehydroemetine and *(-)-S,S*-dehydroisoemetine in *P. falciparum* sensitive strain (3D7A) and two Pf resistant strains (Dd2 and W2). The ratios of *in vitro* cross-resistance of emetine dihydrochloride and *(-)-R,S*-dehydroemetine in both resistant strains (Dd2 and W2) using 3D7A strain as reference were found to be 1.

**Table 1:** **Analysis of results to determine cross-resistance through hypoxanthine incorporation assays**

| Compounds | 3D7A |  | Dd2 |  | W2 |  | ratio<br>Dd2/3D7A | ratio<br>W2/3D7a |
| --- | --- | --- | --- | --- | --- | --- | --- | --- |
| | Pf IC50<br>(μM) | st dev | Pf IC50<br>(μM) | st dev | Pf IC50<br>(μM) | st dev | | |
| Emetine dihydrochloride | 0.234 | 0.022 | 0.269 | 0.025 | 0.181 | 0 | 1 | 1 |
| (-)-*R,S*-dehydroemetine | 0.146 | 0.003 | 0.177 | 0.014 | 0.168 | 0.005 | 1 | 1 |
| (-)-*S,S*-dehydroisoemetine | 1.031 | 0.041 | ND |  | ND |  |  |  |

The results showed that the inhibitory potency observed for emetine dihydrochloride and *(-)-R,S*-dehydroemetine in both multidrug resistant strains (Dd2 and W2) is similar to the sensitive strain 3D7A. These results suggest that there is no cross-resistance with any of the MDR strains tested.

### *In vitro* IC_50_ against *P. falciparum* male and female activated gametes

This bioassay assesses the malaria transmission blocking potential of compounds by estimating their ability to prevent male mature gametocytes to progress to male microgametes or/and to inhibit female gamete activation, as indicators of gametocyte functionality. The activation of male gametocytes into mature microgametes is evaluated by the process of exflagellation (extrusion of rapidly waving flagellum-like microgametes from the infected erythrocyte). The activation of female gametocytes is evaluated based on the specific expression of the Pfs25 protein at the surface of the female activated gametes (45). Male *P. falciparum* gametocytes exflagellate when activated, causing the movement of the surrounding RBCs in the media. Detecting these changes in cells position, we were able to detect activated male gametes. Female *P. falciparum* gametocytes round up when activated and the Pfs25 protein distributes widely in the membrane of the gamete. Using a monoclonal antibody against this protein we were able to specifically detect activated female gametes. Dual Gamete Formation assay was performed and the results are shown in table 2.

**Table 2:**
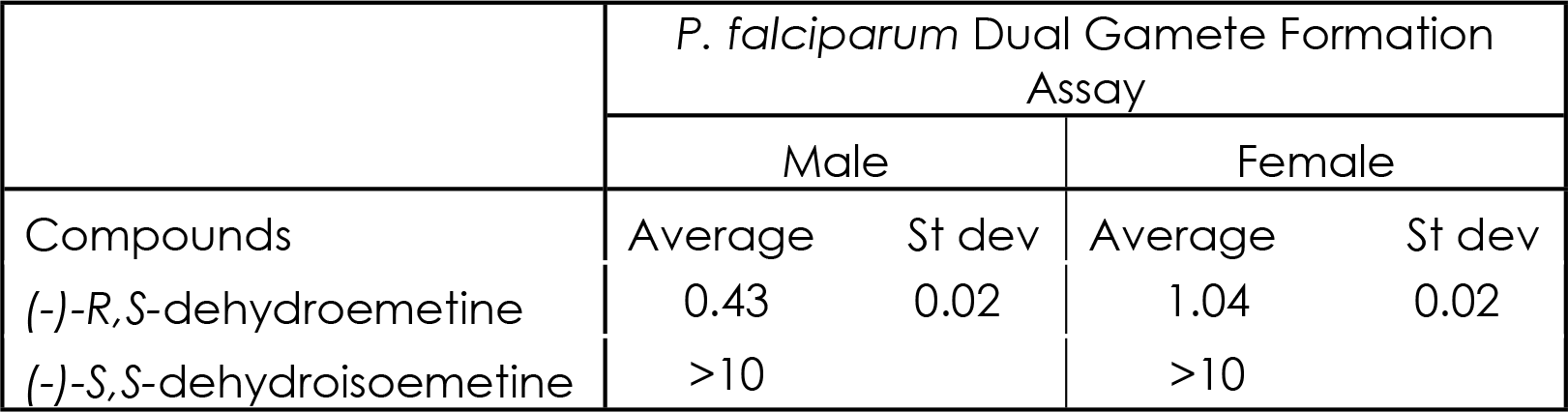
***In-vitro* IC_50_ values of *(-)-R,S*- dehydroemetine and *(-)-S,S*-dehydroisoemetine against male and female gametocytes in *P. falciparum* NF54 strain**

### hERG channel inhibition assay

Dose related cardiovascular side-effects have been observed following treatment of amoebiasis with emetine which included ECG changes such as T-wave inversion, prolongation of QT interval and to some degree increased width of QRS complex and PR interval. Hypotension, tachycardia and precordial pain were also observed (25). Stoppage of treatment resulted in complete recovery of cardiovascular functions. Cardiac microscopic examination revealed a separation of muscle fibres and destruction of myocardial fibres but an absence of inflammatory cells leading to the interpretation that myocarditis is toxic rather than inflammatory in origin. In another study on thirty two patients, pain was noted at injection site along with ECG abnormalities, myalgia, muscle weakness and increased levels of serum creatinine phosphatase (46). In a study on guinea pigs by Schwartz and Herrero (1965), it was observed that dehydroemetine was excreted faster than emetine. It was also postulated that reduced cardiotioxity of dehydroemetine could be due to decreased tissue affinity to the heart in comparison to emetine (25).

In a resting cardiac cell, the concentration of K^+^ is high intracellularly, which creates a chemical gradient for K^+^ to diffuse out of the cells. A subunit of the rapid delayed rectifier potassium ion channel is involved in the cardiac repolarisation (47). It is encoded by the hERG gene (Human ether-a-go-go related gene). Since emetine is known to affect the movements of Na^+^, K^+^ and Ca^2+^ ions, 2,3-dehydroemetine is also thought to affect ion permeability. The study aims to establish if hERG channel inhibition is responsible for the cardiotoxicity observed with emetine therapy in a bid to propose analogues with reduced side effects.

### Data Analysis of hERG channel inhibition assay

For each replicate the hERG response is calculated using the following equation:

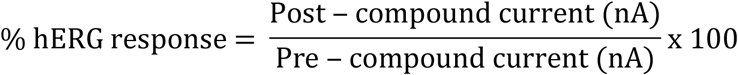

The % hERG response was plotted against concentration for the test compound and, where concentration-dependent inhibition was observed, the data were fitted to the following equation and an IC_50_ value calculated:

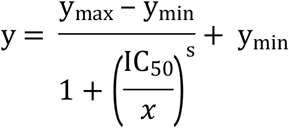

Where: y = hERG response, y_max_ = mean vehicle control response, x = concentration, IC_50_ = concentration required to inhibit current by 50% and s = Hill slope *(-)-R,S*-dehydroemetine has an IC_50_ of 19.3 μM for hERG channel whereas *(-)-S,S*-dehydroisoemetine has an IC_50_ of 2.99 μM. The IC_50_ of quinidine which was used as a positive control is 1.99 μM. Thus it was found that *(-)-R,S*-dehydroemetine is not a potent hERG channel inhibitor, but *(-)-S,S*-dehydroisoemetine is a potent inhibitor.

**Table 3:** **hERG channel inhibition assay results for *(-)-R,S*-dehydroemetine and *(-)-S,S*-dehydroisoemetine**

| Cyprotex | hERG channel inhibition (IC<sub>50</sub> determination) |  |  |  |  |  |  |  |
| --- | --- | --- | --- | --- | --- | --- | --- | --- |
| Compounds | % Mean inhibition |  |  |  |  |  |  |  |
|  | 0<br>μM | 0.016 μM | 0.08<br>μM | 0.4<br>μM | 2 μM | 10<br>μM | 50 μM | IC<sub>50</sub> (μM) |
| (-)-*R,S*-dehydroemetine | 0 | 7.79 | 6.34 | 12.7 | 7.21 | 39.4 | 61.4 | 19.3 |
| (-)-*S,S*-dehydroisoemetine | 0 | 8.13 | 4.61 | 9.33 | 36 | 70.7 | 96.9 | 2.99 |

### Staining with rhodamine123 and draq5 for fluorescence microscopy and measurement of mitochondrial membrane potential

Changes in mitochondrial membrane potential was measured using rhodamine123, a cationic fluorescent dye (48). It was visualised on fluorescence microscopy that rhodamine123 localises with the cytoplasm of the parasites. Draq5 is a far-red fluorescent DNA dye which is cell permeable and was used to identify the parasites. SYBRgreen was not preferred for this experiment as it emits in the same channel as rhodamine123 and would result in the overlap of emission signals.

Changes in mitochondrial membrane potential can be observed using rhodamine123 (excitation wavelength: 511 nm and emission wavelength: 534 nm), a membrane permeable fluorescent dye which accumulates by electrostatic attraction in the mitochondria because of its negative transmembrane potential. A change in the dye’s concentration in the mitochondria is caused by a depolarisation event and can be visualised as a shift in fluorescent intensity of rhodamine123 (49). Draq5, (excitation wavelength: 647 nm and emission wavelength: >665 nm) a DNA dye was used to distinguish the parasites on flow-cytometry in APC-Cy7-A channel. Atovaquone, a known mitochondrial inhibitor was used as a control. A decrease in florescent intensity measured in FITC-A channel indicates a loss of mitochondrial membrane potential. A shift in fluorescent intensity was observed after treatment with all four compounds. Emetine, *(-)-R,S*-dehydroemetine and *(-)-S,S*-dehydroisoemetine showed a shift in fluorescent intensity of rhodamine123 in a direction similar to atovaquone indicating a possible mitochondrial effect (Fig 9).

**Fig 9:**
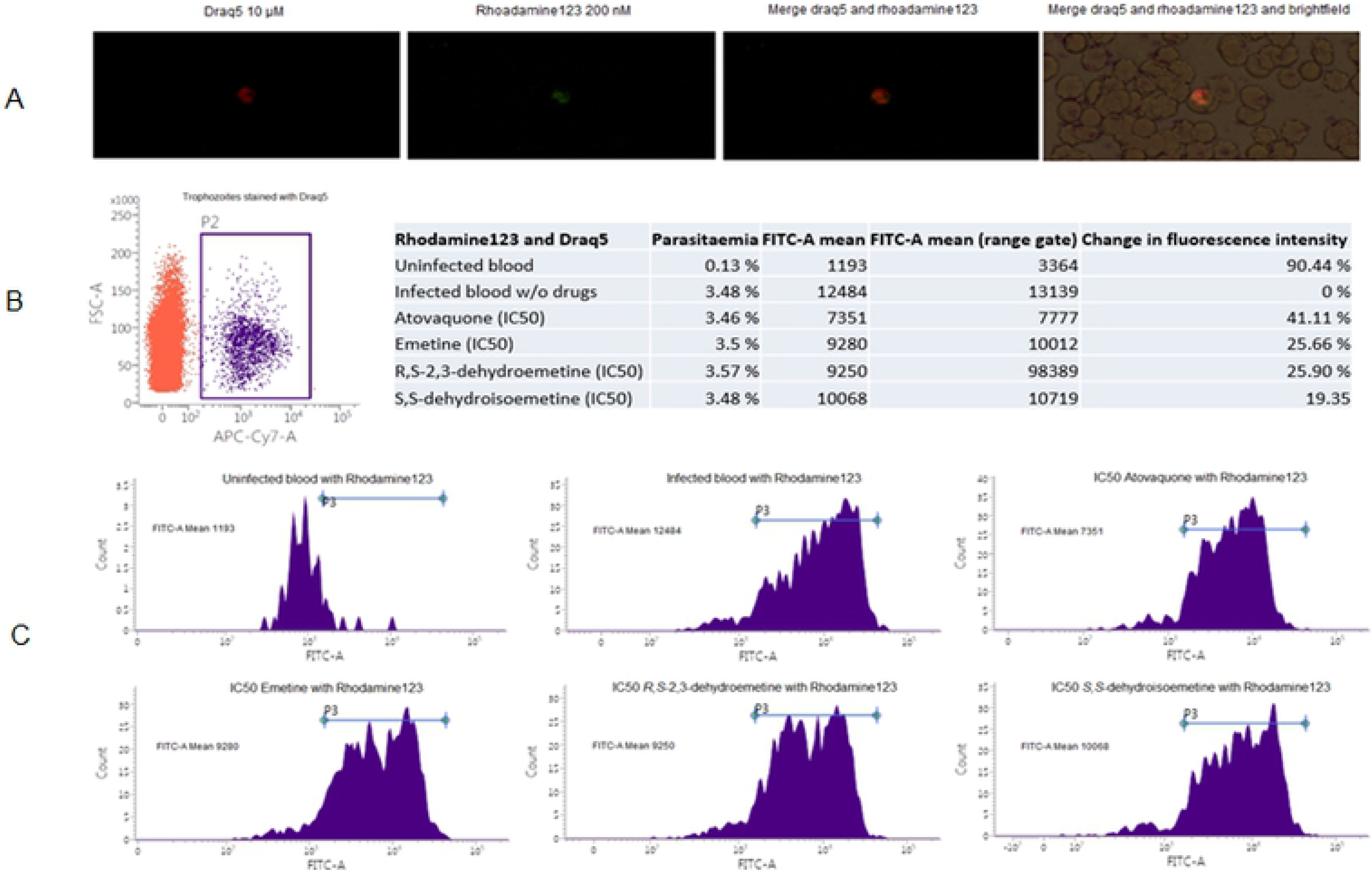
Microscopy and flow-cytometry results for effect of atovaquone, emetine, *(-)-R,S*-dehydroemetine and *(-)-S,S*-dehydroisoemetine on mitochondrial membrane potential. (a) *P. falciparum* K1 strain trophozoites stained with 10 μM draq5 and 200 nM rhodamine123 visualised under fluorescence microscope at 100x magnification, (b) and (c) Changes in mitochondrial membrane potential observed on treatment with IC_50_ of atovaquone, emetine, *(-)-R,S*-dehydroemetine and *(-)-S,S*-dehydroisoemetine

### Validation of the CalcuSyn assay for malaria

CalcuSyn method has been validated in our laboratory as a useful tool anti-malarial drug interaction analysis (38). To establish the robustness of the method the known synergistic drug combination, atovaquone and proguanil, was used. Triplicate samples were analysed by SG-plate-reader after 72 hours. Calclusyn predicted strong synergism between the two drugs which is in accordance with the published literature.

Following validation of the Calcusyn software employing Chou-Talalay method for drug interaction analysis using the atovaquone-proguanil combination, the interaction between *(-)-R,S*-dehydroemetine-atovaquone and *(-)-R,S*-dehydroemetine - proguanil were studied. The doses used for each compound were based on the known ED_50_ values which served as the midpoint for a two-fold, constant-ratio dose series as shown in table 4.

**Table 4:** **The dose series used for the combination of existing antimalarials (AM) with *(-)-R,S*-dehydroemetine**

| Level | (-)-*R,S*-dehydroemetine (nM) | Atovaquone (nM) | Proguanil (uM) |
| --- | --- | --- | --- |
| A | 12.5 | 0.5 | 1.75 |
| B | 25 | 1 | 3.5 |
| C | 50 | 2 | 7 |
| D | 100 | 4 | 14 |
| E | 200 | 8 | 28 |
| F | 400 | 16 | 56 |
| G | 800 | 32 | 112 |
| Combination ratio with (-)-*R,S*-dehydroemetine | -- | 1:25 | 140:1 |

### CalcuSyn-based analysis of the atovaquone - proguanil combination

The Calcusyn based analysis of the drug interactivity between atovaquone and proguanil using a constant-ratio combination of 1:7000 dose was carried out. The output included a dose-effect curve and a median-effect plot in addition to the Combination Index (CI) and an isobologram plot, to figuratively depict the compound’s potency and conformity to the mass action law (Fig 10). Specifically, CI = 0.20, 0.34 and 0.57 at the ED_50_, ED_75_ and ED_90_ levels, respectively were obtained inferring strong synergism at ED_50_ and ED_75_, and synergism at ED_90_. Good correlations coefficients of the median-effect plot were reported for atovaquone (r = 0.99), proguanil (r = 0.89), and the combination (r = 0.93), inferring good conformity to the mass-action law as shown in table 5.

**Table 5:** **CalcuSyn based analysis of drug interaction between atovaquone and proguanil**

| Drug | CI Values at |  |  | Dm | m | r |
| --- | --- | --- | --- | --- | --- | --- |
|  | ED50 | ED75 | ED90 |  |  |  |
| Atovaquone | N/A | N/A | N/A | 6.5262 | 0.88422 | 0.99782 |
| Proguanil | N/A | N/A | N/A | 27771 | 1.06144 | 0.89349 |
| Combination (1:7000) | 0.2088 | 0.34424 | 0.57296 | 0.51518 | 0.68303 | 0.93063 |

**Fig 10:**
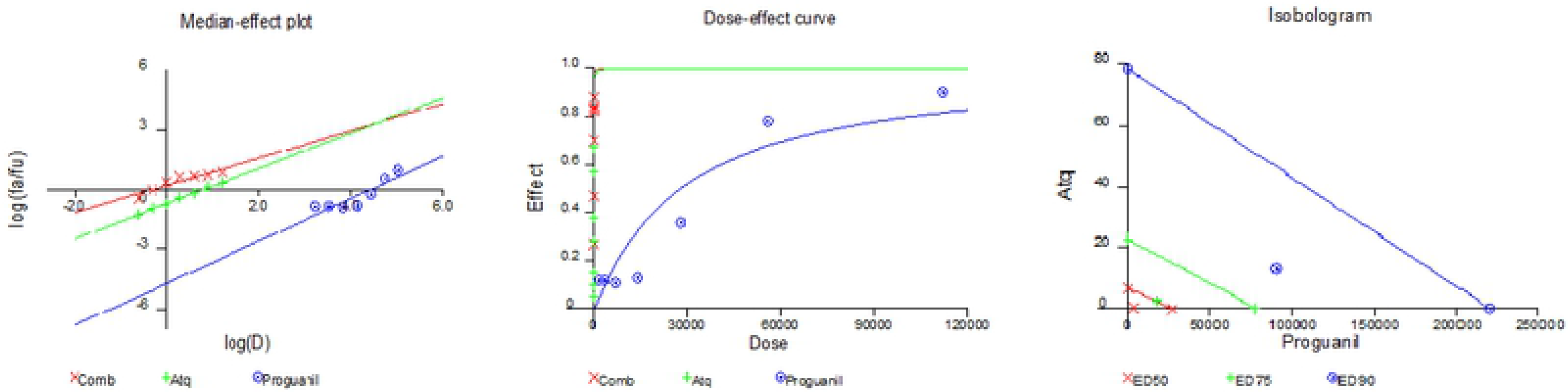
CalcuSyn based median effect plot, dose-effect curve and isobologram for atovaquone and proguanil

### CalcuSyn-based analysis of the (-)-*R*,*S*-dehydroemetine - atovaquone combination

The Calcusyn based analysis of the drug interactivity between (-)-*R*,*S*-dehydroemetine and atovaquone using a constant-ratio combination of 25:1 dose was done (Fig 11). Specifically, CI = 0.88, 0.88 and 0.89 at the ED_50_, ED_75_ and ED_90_ levels, respectively were obtained inferring mild synergism at all measurement points. Good correlations coefficients of the median-effect plot were reported for atovaquone (r = 0.94), (-)-*R*,*S*-dehydroemetine (r = 0.94), and the combination (r = 0.96), inferring good conformity to the mass-action law as shown in table 6.

**Table 6:** **Calcusyn based analysis of drug interaction between atovaquone and (-)-*R,S*-dehydroemetine**

| Drug | CI Values at |  |  | Dm | m | r |
| --- | --- | --- | --- | --- | --- | --- |
|  | ED50 | ED75 | ED90 |  |  |  |
| Atovaquone | N/A | N/A | N/A | 8.56439 | 1.51904 | 0.94197 |
| (-)-R,S-dehydroemetine | N/A | N/A | N/A | 85.98414 | 2.41228 | 0.94623 |
| Combination (1:25) | 0.88215 | 0.88286 | 0.89502 | 2.16472 | 2.08897 | 0.96644 |

**Fig 11:**
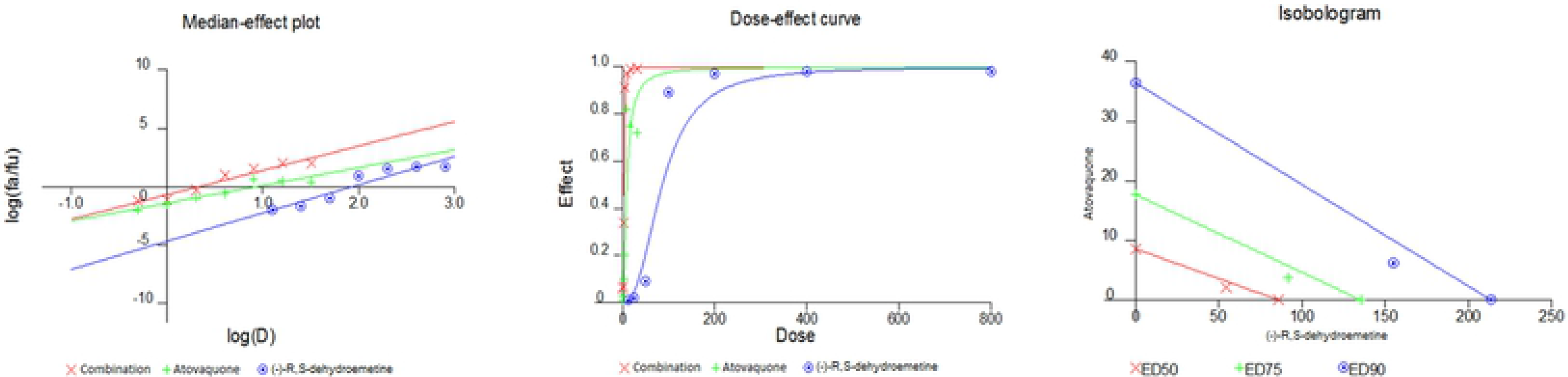
Calcusyn based median effect plot, dose-effect curve and isobologram for atovaquone and *(-)-R,S*-dehydroemetine

### CalcuSyn-based analysis of the *(-)-R,S*-dehydroemetine - proguanil combination

The CalcuSyn based analysis of the drug interactivity between *(-)-R,S*-dehydroemetine and proguanil using a constant-ratio combination of 140:1 dose was done (Fig 12). Specifically, CI = 0.67, 1.04 and 1.62 at the ED_50_, ED_75_ and ED_90_ levels, respectively were obtained implying synergism at ED_50_, near–additive effect at ED_75_ and antagonism at ED_90_. Good correlation coefficients of the median-effect plot were obtained for proguanil (r = 0.90), *(-)-R,S*-dehydroemetine (r = 0.86), and the combination (r = 0.95), suggesting good conformity to the mass-action law as shown in table 7.

**Table 7:** **CalcuSyn based analysis of drug interaction between proguanil and *(-)-R,S*-dehydroemetine**

| Drug | CI Values at |  |  | Dm | m | r |
| --- | --- | --- | --- | --- | --- | --- |
|  | ED50 | ED75 | ED90 |  |  |  |
| Proguanil | N/A | N/A | N/A | 20.931 | 1.61801 | 0.90662 |
| (-)-*R,S*-dehydroemetine | N/A | N/A | N/A | 232.4649 | 0.81711 | 0.86213 |
| Combination ratio (140:1) | 0.67736 | 1.04995 | 1.62774 | 14.16876 | 0.98301 | 0.95879 |

**Fig 12:**
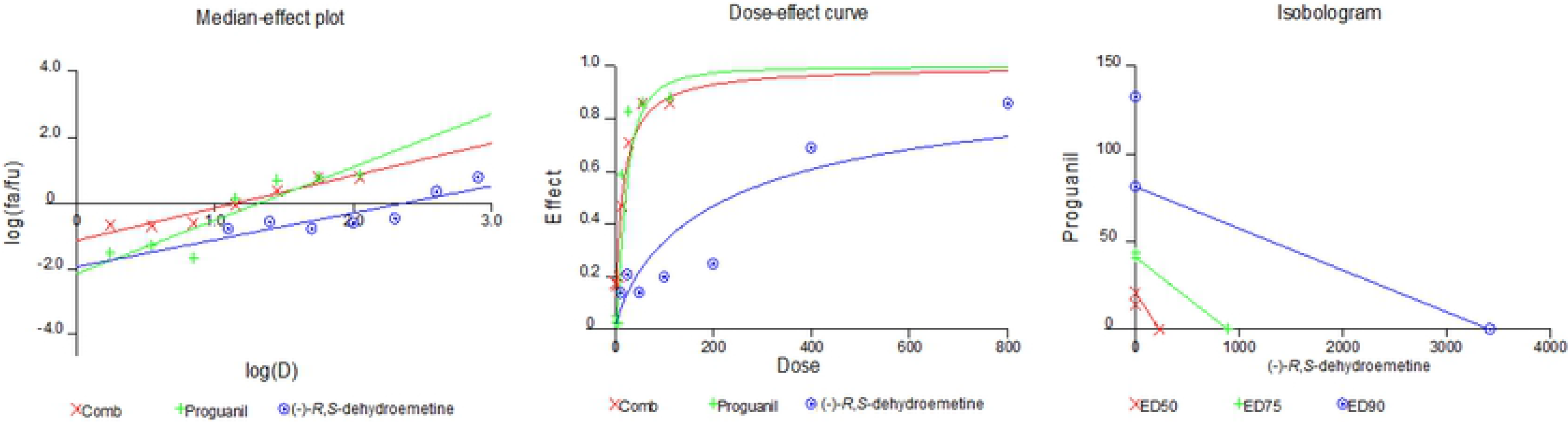
Calcusyn based median effect plot, dose-effect curve and isobologram for proguanil and *(-)-R,S*-dehydroemetine

After 72 hours of incubation, the *(-)-R,S*-dehydroemetine and atovaquone interaction was classified as synergistic at all inhibitory levels analysed. For the *(-)-R,S*-dehydroemetine and proguanil combination the ED_50_ level of inhibition was classified as synergism, the ED_75_ as nearly additive and ED_90_ as antagonism.

## Discussion

In conclusion, we report here the nanomolar antimalarial efficacy of the synthetic emetine analogue*(-)-R,S*-dehydroemetine in the multi drug resistant K1 strain of *Plasmodium falciparum* (IC_50_ of 69.58 ± 2.62 nM). The clinical use of emetine dihydrochloride as an anti-amoebic drug was superseded by the better tolerated synthetic analogue 2,3-dehydroemetine in the 1070s. The isomer *(-)-S,S*-dehydroisoemetine was found to be less potent with an IC_50_ of 1.85 ± 0.2 μM. Molecular modelling suggests that this greater potency of *(-)-R,S*-dehydroemetine is linked to its ability to mimic more fully the interactions of emetine with the E-site of Pf40S subunit, compared to the *(-)-S,S*-dehydroisoemetine isomer. The improvements in the dose-dependent cardiotoxicity previously reported for 2,3-dehydroemetine in comparison to emetine dihydrochloride was linked to decreased affinity to cardiac myocytes and increased clearance from the body (27). We propose that the very significant differential in vitro activities of emetine dihydrochloride in *Entamoeba histolytica* (IC_50_ 26.8 ± 1.27 μM) and *Plasmodium falciparu*m (IC_50_ 47 ±; 2.1 nM) (13) (14) would mean that the dose related toxicity profile for its re-positioned use will be different.

Analysis done by GSK showed that in asexual blood stages, the compounds exhibited no cross-resistance issues. Mosquito vector dynamics, the number of people with peripheral gametocytemia in the population, and the infectiousness of circulating gametocytes to mosquitoes determines the transmission of malaria. Interrupting transmission is an important aspect of preventing malaria in endemic areas and there has been renewed interest in compounds preventing the formation of gametocytes. *(-)-R,S*-Dehydroemetine was found to be gametocidal. It displayed activity against both male and female gametes. Thus it also has the potential to block the transmission of malaria.

The selectivity indices calculated using MTT assays in HepG2 cell lines was similar for emetine and its analogues. It is important to note that the low selectivity index was expected given the documented anticancer properties of emetine (50). This could explain the low IC_50_ obtained on Hep G2 lines as these are rapidly multiplying cancer cell lines. The selective advantage of *(-)-R,S*-dehydroemetine for cardiotoxic effects is linked to its faster elimination from body, hence the appropriate toxicity investigation to define this would be the use of in vivo animal models. It is imperative to progress this work in this direction. It was also clearly established that the cardiac toxicity for *(-)-R,S*-dehydroemetine was not effected through the hERG channel. Our observations suggest that emetine and its two synthetic analogues have atovaquone-like activity upon measurement of changes in the mitochondrial membrane potential at IC_50_ concentrations, indicating multi-modal mechanisms of action. The failure of monotherapy has been unambiguously demonstrated in malaria, hence, the forceful insistence by the WHO for the use of artemisinin-based combination therapy (ACT) as a policy standard. In addition to more potent therapeutic efficacies, other benefits of combination regimes include decreased toxicity, favourable synergistic interactions and most significantly, the potential to impede or delay the onset of resistance. Drug interaction analyses presented here emphasise a route to further dose reduction to minimise toxicity. *(-)-R,S*-Dehydroemetine was found to exhibit synergistic activity with atovaquone and proguanil.

Emetine and its analogues, constitute a pleiotropic group of natural product-derived compounds which could potentially enhance the depleted antimalarial armamentarium, should a catastrophic gap in the drug market occur as a result of the spread of resistance to frontline anti-malarials. The work presented here provides strong justification for further optimisation, particularly for use in a hospital setting in cerebral malaria, where appropriate monitoring could obviate progression of the reversible cardiotoxicity previously reported. Its hepatic concentration ability could indeed have favourable consequences for the treatment of Plasmodium vivax infections. The history of antimalarial chemotherapy has been largely reliant on natural product-derived leads. We provide here a significant body of evidence to progress and further optimised a previously overlooked, potent and affordable natural product compound.

## Materials and Methods

### Culture of *Plasmodium falciparum*

RPMI 1640 media containing 25 mM HEPES and 0.3 g/L L-glutamine (Gibco, Life Technologies, UK) supplemented with sterile filtered 2.5 g Albumax (Sigma, UK), 2.5 ml hypoxanthine (Sigma, UK), 2.5 ml 40% glucose (Dextrose Anhydrous, Fisher Scientific, UK) and 0.5 ml gentamycin (Sigma, UK), were used for the culture of erythrocytic stage, strain K1, *P. falciparum* parasites (gifted by Prof John Hyde, University of Manchester, UK, original source: Thai-K1 clone), under a 5% CO_2_ 5% O_2_ and 90% N_2_ gas mixture (BOC Limited, UK) at 37°C. All routine culture methods were consistent with those employed by (51).

O+ human blood (purchased from NHS Blood Bank, Manchester, UK) was used routinely to maintain the parasites. Continuous cultures were maintained at 5% haematocrit. Since synchronous parasite development is observed in natural hosts and this synchrony is lost quite rapidly in *in-vitro* cultures, sorbitol was used to keep the parasites in tight synchrony. In brief, the parasitized blood pellet was re-suspended 1:10 in 5 % sorbitol (prepared in distilled water and filtered using 0.22 μm porosity Millipore filter), incubated at room temperature for 5 minutes and then centrifuged at 3000 rpm for 5 minutes. The supernatant was removed and complete media was used to wash the pellet 3 times before setting up a new culture.

### Synthesis of (-)-2,3-Dehydroemetine

(-)-2,3-Dehydroemetine was synthesised as per the established literature (Brossi 1967, Patent No. 3,311633 [16]). The synthesis was outsourced to Chiroblock GMBH, Germany. Previously published patent literature by chemists at Hoffmann-La Roche did not include modern analytical data such as mass spectrometry or NMR. The methodology adopted for the synthesis *(-)-R,S*-dehydroemetine **7** used in this study followed the published patent with minor modifications as depicted in (see Fig 13) below.

**Fig 13:**
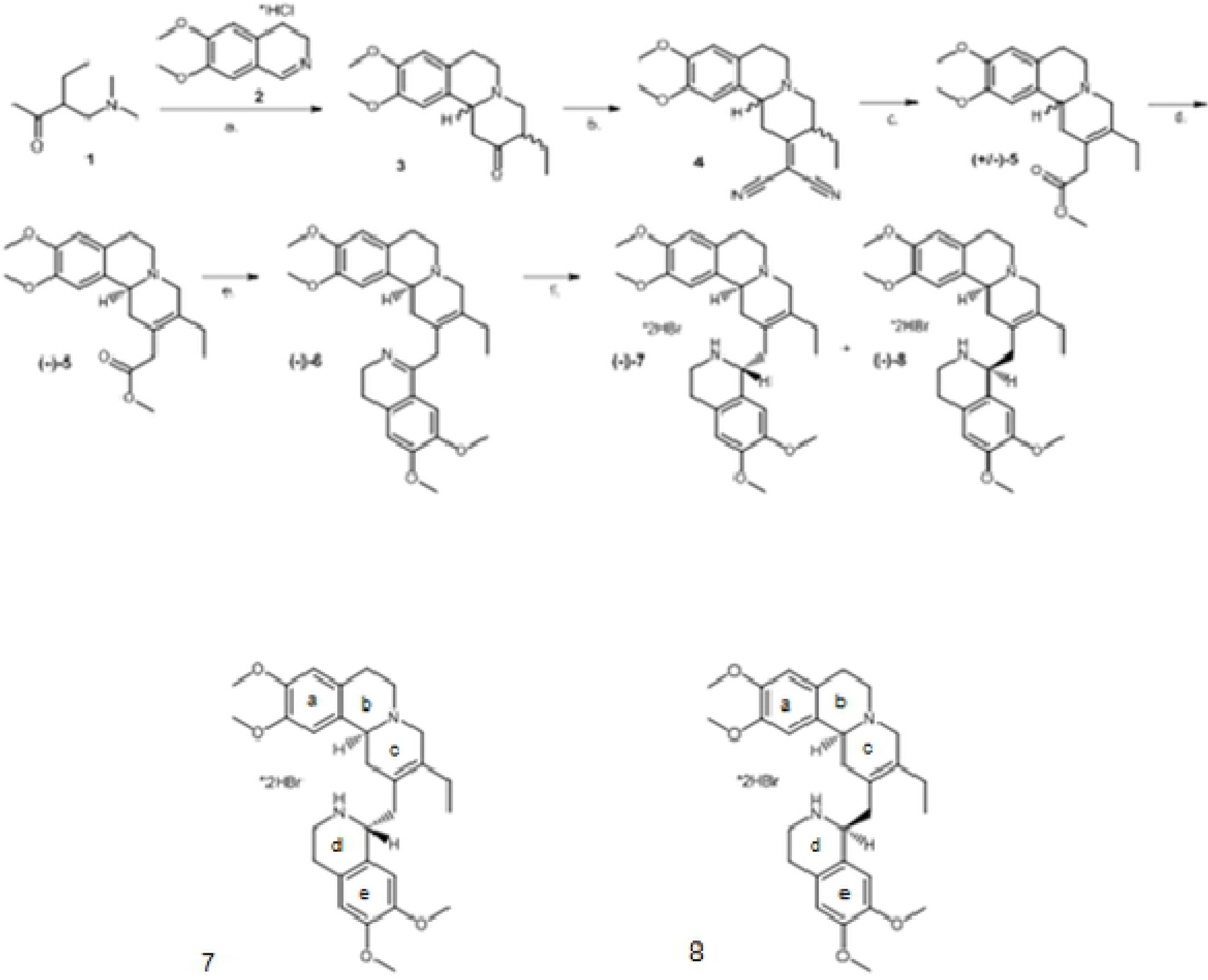
Synthesis of 2, 3-dehydroemetine. Conditions: a) i. iodomethane, ethanol, rt, 24 h, ii. **2**, KOAc, reflux, 3 h, 72%; b) malononitrile, ammonium acetate/acetic acid, toluene, reflux, 2.5 h, 69%; c) i. 20% HCl, reflux, 5 h, ii. methanolic HCl, rt, 18 h, 45%; d. (+)-dibenzoyl tartrate, methanol, two recrystallizations, 25%; e) i. 3M HCl, reflux, 90 min, ii. 3,4-dimethoxyphenethylamine, xylenes, reflux, 18 h, iii. POCl_3_, benzene, reflux, 1 h, 28%; f) NaBH_4_, methanol, rt, 1 h, 81% of a 1:1 diastereomeric mixture. 7: (*S*)-2-((*R*)-6,7-Dimethoxy-1,2,3,4-tetrahydro-isoquinolin-1-ylmethyl)-3-ethyl-9,10-dimethoxy-1,6,7,11b-tetrahydro-4H-pyrido[2,1-a]isoquinoline * dihydrobromide 8: (S)-2-((S)-6,7-Dimethoxy-1,2,3,4-tetrahydro-isoquinolin-1-ylmethyl)-3-ethyl-9,10-dimethoxy-1,6,7,11b-tetrahydro-4H-pyrido[2,1-a]isoquinoline * dihydrobromide

The two compounds synthesised were (-)-*R,S*-dehydroemetine, **7**, and (-)-*S,S*-dehydroisoemetine, **8**. Briefly, the Mannich base **1** was quaternized and reacted with the imine **2** giving piperidinone **3** as an approximately 5:1 mixture of diastereomers which were not separated. Knoevenagel condensation with malononitrile led to unsaturated dinitrile **4** in 69% yield. Hydrolysis and decarboxylation, with concomitant alekene isomerization, gave **5** after methylation. Resolution with (+)-dibenzoyl-*D*-tartaric acid gave homochiral (-)-**5**. Hydrolysis and amidation with homoveratrylamine followed by Bischler–Napieralski cyclization led to dihydroisoquinoline derivative **6**, also known as 2-dehydro-*O*-methyl psychotine. The cyclic imine was reduced with sodium borohydride in methanol to give a 1:1 mixture of the desired 2,3-dehydroemetine **7** and 2-dehydroisoemetine dihydrobromide **8**, separated by fractional crystallization. The purity of (-)-*R,S*-dehydroemetine obtained through this method was 85% with the balance consisting of the diastereomer, **8**.

### Molecular modelling

In preparation for predicting the bound pose of emetine, *(-)-R,S*-dehydroemetine and *(-)-S,S*-dehydroisoemetine, the cryo-EM structure of *P. falciparum* 80S ribosome bound to emetine dihydrochloride was obtained from the Protein Data Bank (PDB code 3J7A) (16). We note that this cryo-EM structure features the enantiomer of emetine rather than emetine itself; therefore, an initial 3D structure of emetine was obtained from the OMEGA-generated conformer in the Pubchem database (55). Subsequently emetine derivatives *(-)-R,S*-dehydroemetine and *(-)-S,S*-dehydroisoemetine were constructed using the software package MOE (56). Prior to *in silico* docking, crystallographic water molecules and a magnesium ion were removed (the latter was within 13 Å of the active site and was found to unduly influence docking solutions). Hydrogens were assigned consistent with physiological pH. Docking of ligands to the *Pf*40S subunit was performed using MOE-Dock[Chemical Computing Group Inc., 1010 Sherbrooke St. West, Suite #910, Montreal, QC, Canada, H3A 2R7, 2016]. The binding site region was defined using the cryo-EM ligand location. Ligand placement used the Triangle Matcher protocol. Internal flexibility was permitted for ligands but not ribosome. Poses were scored via the London dG scoring function. While this docking protocol was able to reproduce well the cryo-EM pose of the emetine stereoisomer bound to the Pf40S subunit (S1 Figure, Supporting Information), please see comment later regarding this ligand’s structure.

### Drug preparation

Emetine dihydrochloride was obtained from Sigma Aldrich, UK. The dehydroemetine analogues were synthesised as per the established literature (Scheme 1, Brossi 1967, Patent No. 3,311633 [16]). Derivatives were validated using NMR, HPLC and FTIR. The synthesis process was outsourced to Chiroblock GMBH, Germany. Drug stock solutions were prepared in DMSO in accordance with manufacturer instructions and primary stock concentration of 5 mM was aliquoted and stored at −20°C until further use. Serial dilutions of the working solution were made using complete media for the experiments.

### Drug dosing for IC_50_ determination

Refined dose ranges were selected for emetine dihydrochloride (12.5 nM – 200 nM), *(-)-R,S*-dehydroemetine (12.5 nM - 200 nM), and *(-)-S,S*-dehydroisoemetine (0.625 μM - 10 μM) to permit accurate IC_50_ calculation against the K1 strain of *P. falciparum*. Ring stage parasites were diluted to 0.5-1% parasitaemia, 2.5% haematocrit (in a 96 well plate format, 200 μl final well volume) and treated for 72 hours. Dose response parasitaemia was determined using the SYBR Green flow cytometer method previously optimised in the laboratory (14).

### SYBR Green staining of erythrocytic stage *P. falciparum* for flow cytometry

Following drug efficacy experiments, 150 μl from the control and drug-treated wells on a 96 well plate were transferred to a microcentrifuge tube and samples were washed once with PBS. The supernatant was removed and samples were incubated in the dark at room temperature for 20 minutes following addition of 1 ml of 5 × SYBR Green solution (prepared by adding 5 μl of 10,000 × SG to 10 ml PBS). After staining, the samples were centrifuged for 1 minute at 14,000 rpm and the supernatant was discarded. Samples were re-suspended in 250 μl of 0.37% formaldehyde fixation solution prepared by diluting 36.5% formaldehyde (Sigma, UK) with PBS to a final concentration of 0.37%. The samples were placed in the fridge and incubated at 4℃ for 15 minutes. Subsequently, PBS was used to wash the samples 3 times and finally the samples were suspended in 1 ml PBS. Parasitaemia was determined by SYBR green fluorescence using the FITC channel of the BD FACsVerse flow cytometer system (Blue laser, excitation laser line 488 nm EX_max_ 494 nm/Em_max_ 520 nM) and cell size (forward scatter, FSC-A). Fifty thousand events were recorded and in three replicates for each. Fluorescent events in drug-treated samples were compared with infected and uninfected blood counterparts and gated accordingly to obtain the percentage parasitaemia.

### IC_50_ speed assay

The assay was used to determine the speed of action of the emetine analogues. Unsynchronised cultures of K1 strain *P. falciparum* was used and parasites were grown in presence of *(-)-R,S*-dehydroemetine and *(-)-S,S*-dehydroisoemetine for three incubation periods of 24 hours, 48 hours and 72 hours. The assays were analysed by determining SYBR green fluorescence as described above in the methodology.

### Stage-specificity assay

Parasite cultures with ≥ 80% trophozoites and ≥ 80% rings were obtained by synchronisation with 5% Sorbitol (42). The cultures were synchronised twice, at 0 hours and 31 hours to obtain young rings which were up to 3 hours old. To obtain early schizont stages, the culture was synchronised twice with the second synchronisation being 6 to 8 hours after the first. Each synchronous stage was incubated for 24 hours at 37 °C on a 96-well microtitre plate with a two-fold serial dilution of the drugs ranging from 1.6 to 100 fold the IC_50_ of each drug. The plates were washed 4 times after the incubation in order to dilute the drug concentration by > 1000 fold. The plates were incubated for another 24 hours at 37°C following which SYBR green staining was used to read the plates as described above.

### MTT Assay for testing cell cytotoxicity

HepG2 cells (University of Salford stocks purchased from American Type Culture Collection (ATCC), USA), were grown in media consisting RPMI 1640, 2 mM *L*-Glutamine, HEPES, 10% foetal bovine serum, 1% non-essential amino acids and 1% penicillin-streptomycin. Cells were calculated on a haemocytometer to make a new cell-media solution to a concentration of 5000 cells/100 μl and 100 μl of this new cell solution was added into each well of the 96 well plate. After 24 hours incubation, 100 μl of the drug prepared in the media were added to the wells. This was repeated in triplicate and the plate was incubated for a further 24 hours (52). After the incubation period, 50 μl of MTT solution was added to each well and incubated for 3 hours. The liquid in each well was aspirated carefully and 200 μl of DMSO was added to each well and results were read on the Ascent plate reader.

### Derivation of dose-response curves and IC_50_ values

The infected blood controls were set at 100% and percentage parasitaemia was calculated for drug treated samples relative to the infected control. For IC_50_, GraphPad prism 5.0 was used to further process the data. The largest value in the data set was set to correspond to 100% and the smallest value to 0% by normalising the data. IC_50_ values were calculated using nonlinear regression (Graphpad prism 5.0) by using log-transformed drug concentrations plotted against the dose response. The log(inhibitor) vs. Normalised response-Variable slope option was used for IC_50_ calculation. Emetine dihydrochloride was used as a control drug to validate the method.

### Determination of cross-resistance through IC_50_ determination in multidrug resistant *P. falciparum* strains (carried out at Tres Cantos, GSK)

The three strains used for the assay were 3D7A, Dd2 and W2 based on their resistance profiles. 3D7A is chloroquine–sensitive, Dd2 is resistant to chloroquine, mefloquin and pyrimethamine (53) and W2 is resistant to chloroquine, quinine, pyrimethamine, cycloguanil, and sulfadoxine (54) (from the Malaria Research and Reference Reagent Resource Center MR4). A culture of parasitized red blood cells (RBC) of the corresponding strain, (0.5% parasitemia, 2% haematocrit) in RPMI-1640, 5% albumax and 5 μM hypoxanthine was exposed to 9 dilutions (3-fold serial dilutions) of the compound starting at 5 μM. 100 μl of culture volume was plated in 96-well flat bottom microtiter plates + 0.5 μl drug (stock x200 in DMSO). Plates were incubated for 24 h at 37 °C, 5% CO2, 5% O_2_, 90% N_2_. Next, 3H-hypoxanthine was added and plates were incubated for another 24 h period. After that, parasites were harvested on a glass fiber filter using a TOMTEC Cell harvester 96. Filters were dried and melt-on scintillator sheets were used to determine the incorporation of 3Hhypoxanthine. Radioactivity was measured using a microbeta counter. Data are normalized using the incorporation of the positive control, (parasitized red blood cells without drug). IC_50_ values were determined using Grafit 7 program (41).

### Determination of transmission blocking potential through *in vitro* inhibition of gamete activation

Asexual cultures of *P. falciparum* NF54 strain parasites were used to seed gametocyte cultures at 0.5% parasitemia, 4% hematocrit in 50 ml total volume under 3% O_2_/5% CO_2_/92% N_2_ gas. Culture medium (RPMI 25 mM HEPES, 50 mg/l hypoxanthine, 2 g/l NaHCO3 without *L*-glutamine + 5% human serum and 5% Albumax) was replaced daily for 14 days. At day 14, the concentration of non-purified cultures was adjusted to plate 700.000 total cells per well in each 384-well plate. The test drugs were then added in two-fold serial dilutions starting at 10 μM (10 μM – 0.00976 μM) and incubated 48 h at 37 °C (3% 0_2_, 5% CO_2_, 92% N_2_). DMSO was used as the negative control and Thiostrepton as the positive control. Activation was performed with ookinete medium (Same RPMI base used for culture but supplemented with xanthurenic acid 50 μM) supplemented with the antibody anti-Pfs25-Cy3 at a final concentration of 1/2000 (from 1 mg/ml stock). Plates were analyzed to detect exflagellation centres. "Triggered" cultures were then incubated (protected from light) at 26 °C for 24 h (in a thermoregulated incubator). Then, plates were analyzed to detect female activated gametes. Activation of male gametes was detected based on light changes provoked by flagella movements which caused movement of surrounding cells. A 10-frame video was taken and then analyzed to determine these changes in cell position based on pixels change. Then, the script determined where exflagellation centers were located, based also on size and intensity of light changes. Activation of female gametes was based on detection of fluorescent Cy3-Anti Pfs25 antibody (as the primary parameter) followed by a selection of events according to their size, roundness and the intensity of the fluorescence. Both measurements were performed using an automated inverted microscope Ti-E Nikon using JOBS software. Analysis of images and videos was performed with the ICY program. The IC_50_ value was determined using Microsoft Excel and Graphpad.

### hERG Channel Inhibition (IC_50_ Determination) assay Protocol

The experiment to test the potential of emetine, *(-)-R,S*-dehydroemetine and *(-)-S,S*-dehydroisoemetine to inhibit hERG channel was outsourced to Cyprotex, UK. 100 μl of a 20 mM concentration of all three compounds in DMSO was provided to Cyprotex. Compound dilutions were prepared by diluting a DMSO solution (default 10 mM) of the test compound using a factor 5 dilution scheme into DMSO, followed by dilution into extracellular buffer such that the final concentrations tested are typically 0.008, 0.04, 0.2, 1, 5, 25 μM (final DMSO concentration 0.25%). Chinese Hamster Ovary cells expressing the hERG potassium channel were dispensed into 384-well planar arrays and hERG tail-currents were measured by whole-cell voltage-clamping. Amphotericin B was used as a perforating agent is circulated underneath the PatchPlate™ to gain electrical access to the cell. The pre-compound hERG current was measured. Emetine, *(-)-R,S*-dehydroemetine and *(-)-S,S*-dehydroisoemetine in a range of concentrations were then added to the cells and a second recording of the hERG current was made. The test compound was left in contact with the cells for 300 sec before recording currents. Quinidine, an established hERG inhibitor, was included as a positive control, and vehicle control (0.25% DMSO) as the negative control. All buffers, cell suspensions and drug compound solutions were at room temperature. The percent change in hERG current was measured and used to calculate an IC_50_ value.

Each concentration was tested in 4 replicate wells on the PatchPlate™ (maximum of 24 data points). It was ensured that only acceptable cells were used to assess hERG inhibition by applying filters. The cell must maintain a seal resistance of greater than 50 MOhm and a pre-compound current of at least 0.1 nA, and ensure cell stability between pre-and post-compound measurements.

### Staining with rhodamine123 and draq5 for fluorescence microscopy and mitochondrial membrane potential disruption

Rhodamine123 and draq5 were used to observe the effect of emetine and its two synthetic analogues on mitochondrial membrane potential. Rhodamine123 is a mitochondrial specific dye which emits in FITC channel on flow-cytometry whereas draq5 stains the DNA and emits in APC-Cy7-A channel. The two dyes were chosen to perform the experiment as there is very minimal overlap in their emission signals. Synchronised culture of *P. falciparum* K1 strain at trophozoite stage was incubated with 1 ml of 200 nM Rhodamine123 for 1 hour, followed by a further incubation with 100 μl of 10 μM Draq5 for 20 mins. Smears prepared after the incubation were immediately viewed under the fluorescence microscope at 100 x magnification to visualise localisation of the dyes inside the parasite.

To test the effect of emetine and its analogues on mitochondria, synchronised cultures of *P. falciparum* K1 strain at trophozoite stage were incubated at 2.5% haematocrit (in a 96 well plate format, 200 μl final well volume) for 2 hours with IC_50_ concentrations of atovaquone, emetine hydrochloride, *(-)-R,S*-dehydroemetine and *(-)-S,S*-dehydroisoemetine. Compounds were then washed away by centrifuging the cultures and a pellet was prepared for each drug concentration. Pellets were then incubated with 1 ml of 200 nM rhodamine123 for 1 hour, followed by a further incubation with 100 μl of 5 μM draq5 for 20 mins. After a wash with PBS, the experiment was read with flow-cytometer on FITC channel. Loss of mitochondrial membrane potential was indicated by a decrease in fluorescent intensity.

### Drug interaction analysis for *(-)-R,S-dehydroemetine*

Primary stock solutions were prepared as mentioned above. For experimental set up the primary stock solutions were further diluted with complete medium to give final test concentrations. A dose range of 0.125–8 × the ED_50_ was made by a two-fold serial dilution for atovaquone, proguanil, and *(-)-R,S*-dehydroemetine. For atovaquone, the doses ranged from 0.25nM to 16nM for combination with proguanil, and 0.5 nM to 32 nM for combination with *(-)-R,S*-dehydroemetine. For proguanil the doses ranged from 1.75 μM to 112 μM. For *(-)-R,S*-dehydroemetine the doses ranged from 12.5 nM to 800 nM. At each level, the compounds were co-administered for example ED_50_ of atovaquone was combined with the ED_50_ of *(-)-R,S*-dehydroemetine and 2 × ED_50_ atovaquone was combined with 2 × ED_50_ *(-)-R,S*-dehydroemetine and so forth. Parasites were treated at ring stage and incubated for 72 hours in a 96 well plate format. The SYBR Green Plate reader method was used to determine drug susceptibility. The data was analysed for the median-effect using CalcuSyn software (Biosoft) by converting triplicate data to an averaged percentage. The CalcuSyn software generates the CI over a range of f_a_ levels at different growth inhibition percentages. The interpretation of CI was done in accordance with table 8 (37).

**Table 8:** **Classification of synergism or antagonism using CI values generated by Chou-Talalay method (Source: Calcusyn manual, Biosoft, 2006)**

| Range of CI | Symbol | Description |
| --- | --- | --- |
| <0.1 | +++++ | Very strong synergism |
| 0.1-0.3 | ++++ | Strong synergism |
| 0.3-0.7 | +++ | Synergism |
| 0.7-0.85 | ++ | Moderate synergism |
| 0.85-0.90 | + | Slight synergism |
| 0.90-1.10 | ± | Nearly additive |
| 1.10-1.20 | - | Slight antagonism |
| 1.20-1.45 | -- | Moderate antagonism |
| 1.45-3.3 | --- | Antagonism |
| 3.3-10 | ---- | Strong antagonism |
| >10 | ----- | Very strong antagonism |

### Ethics Statement

For routine malaria culture, anonymised whole blood packs deemed unfit/outdated for clinical use were purchased from the NHS Blood Bank at Plymouth Grove, Manchester, UK. For experiments carried out in GlaxoSmithKline, Diseases of the Developing World Medicines Development Campus, Tres Cantos, Spain, the human biological samples were sourced ethically and their research use was in accord with the terms of the informed consents under an IRB/EC approved protocol.

## Acknowledgement

We would like to acknowledge the receipt of Pump Priming Award from the University of Salford, Manchester.

## Supporting Information

**S1 Figure:**
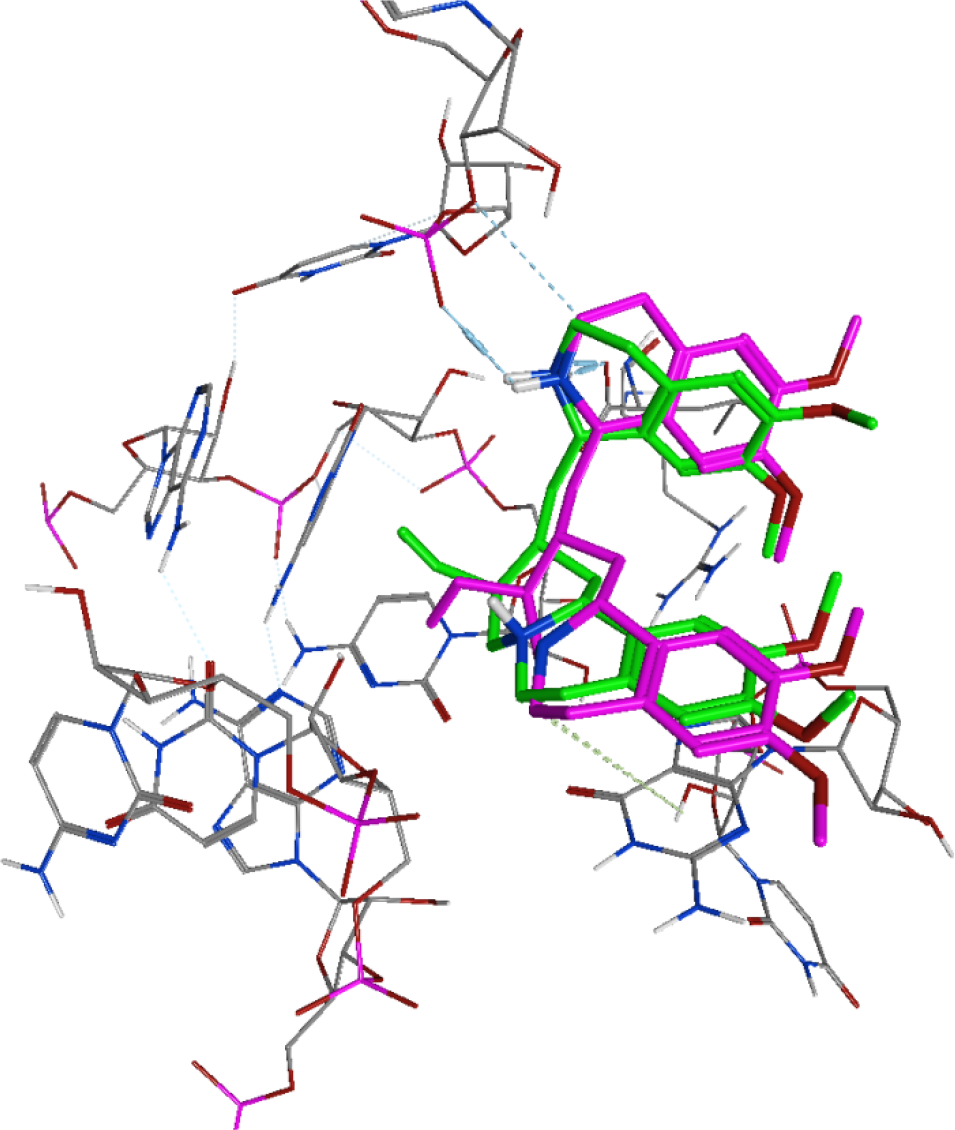
Overlay of docked pose of emetine (pink) and its cryo-EM geometry (green).

